# Quantifying configuration-sampling error in Langevin simulations of complex molecular systems

**DOI:** 10.1101/266619

**Authors:** Josh Fass, David A. Sivak, Gavin E. Crooks, Kyle A. Beauchamp, Benedict Leimkuhler, John D. Chodera

## Abstract

While Langevin integrators are popular in the study of equilibrium properties of complex systems, it is challenging to estimate the timestep-induced discretization error: the degree to which the sampled phase-space or configuration-space probability density departs from the desired target density due to the use of a finite integration timestep. In [1], Sivak *et al*. introduced a convenient approach to approximating a natural measure of error between the sampled density and the target equilibrium density, the KL divergence, in *phase space*, but did not specifically address the issue of *configuration-space properties*, which are much more commonly of interest in molecular simulations. Here, we introduce a variant of this near-equilibrium estimator capable of measuring the error in the configuration-space marginal density, validating it against a complex but exact nested Monte Carlo estimator to show that it reproduces the KL divergence with high fidelity. To illustrate its utility, we employ this new near-equilibrium estimator to assess a claim that a recently proposed Langevin integrator introduces extremely small configuration-space density errors up to the stability limit at no extra computational expense. Finally, we show how this approach to quantifying sampling bias can be applied to a wide variety of stochastic integrators by following a straightforward procedure to compute the appropriate shadow work, and describe how it can be extended to quantify the error in arbitrary marginal or conditional distributions of interest.

## 1. Introduction

Langevin dynamics [2] is a system of stochastic differential equations which describes the behavior of condensed phase systems subject to random weak collisions with fictitious bath particles at thermal equilibrium. In this article we are concerned with the efficient numerical simulation of the Langevin dynamics system. The equations governing the *i*th atom of an *N*-body Langevin system are

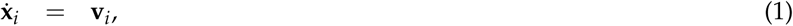

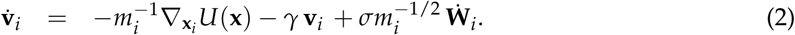

Here, **x***_i_* and **v***_i_* denote the position vector and velocity vector of the *i*th particle of the system (typically each is a vector in ℜ^3^), *m_i_* is the particle mass, and 𝒰(***x***) is the total potential energy, assumed to be a function of all the coordinates, **x** = (**x**_1_,**x**_2_, …,**x**_*N*_). The constant *σ*^2^ = 2*k_B_*T_*γ*_ quantifies the rate of heat exchange with the bath, where *k_B_*T denotes the thermal energy per degree of freedom, *γ* is the collision rate (with dimensions of inverse time), and **W***_i_*(*t*) is a standard 3-dimensional Wiener process [3,4]. It is easily shown that Langevin dynamics preserves the canonical distribution with stationary density

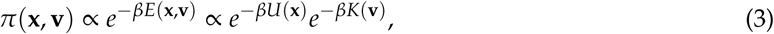

where *β* ≡ (*k*_B_*T*)^−1^ is the inverse temperature, *E*(**x,v**) is a separable energy function *E*(**x,v**) = 𝒰(**x**) + *K*(**v**), and *K*(**v**) = **v**^*T*^*M***v**/2 is the kinetic energy in which *M* is a diagonal matrix constructed from the masses. We assume that the system (1)-(2) is ergodic, meaning that the canonical density is the unique stationary distribution; almost all stochastic paths consistent with (1)-(2) will sample the canonical distribution with density (3).

On a computer, approximating the solution of equations (1)-(2) requires discretizing these equations in time to produce a *finite-timestep Langevin integrator* which can be iterated to compute equilibrium or dynamical properties [5]. A wide variety of schemes have been proposed for this discretization [6–15]. For compact presentation of integration methods, we recast the equations (1)-(2) in vectorial form:

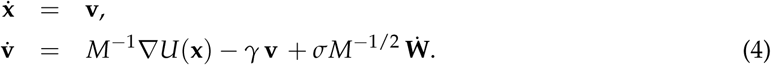

Popular numerical methods for integrating (4) can then be viewed as defining Markov chains on the phase space (**x,v**), with (**x**_*k*+1_, **v**_*k*+1_) defined in relation to (**x***_k_*, **v***_k_*) where the subindex *k* from this point forward should be taken to be a timestep index. Assuming ergodicity, these schemes provide a practical means of sampling the equilibrium distribution.

Until now, assessing whether specific integrators sample the true equilibrium density with greater fidelity than others in specific settings has relied on computing low-dimensional marginal distributions of specific observables perceived to be sensitive to configuration-space sampling errors [16], such as radial distribution functions, marginal distributions of internal coordinates [17], or the configurational temperature [14,18,19]. While it is clear that some observables are more sensitive to errors in configuration-space density than others (Figure 3), and the error in the observables of interest is paramount for a particular application, this highlights the risk of using the error in a single physical property as a surrogate for judging integrator quality, as the error in other observables of interest may be large despite small error in the test observable.

To evaluate numerical Langevin integrators, there would be great utility in a *computable, universal measure* of the bias they introduce in specific concrete settings, such that low error in this measure ensures low error in all observables of interest. There is currently no computable measure of the total configuration-sampling bias for concrete choices of integrator parameters and target system.

Controlling the magnitude of the integrator-induced sampling bias is crucial when computing quantitative predictions from simulation. Following [20], we assume that the numerical method samples an associated probability density *ρ*(**x, v**) which differs from the exact canonical probability density *π*(**x,v**) of (3). Because *ρ* does not have a closed-form, easily computable expression^1^, it is difficult to quantify the error introduced by a given choice of timestep or integrator. We will show how, for a particularly useful measure of error, we can circumvent this problem and develop a simple, effective approach to measuring error in complex molecular systems. Our focus in the sequel is on the error in configurational averages. We therefore introduce the notation *ρ*_**x**_ and *π*_**x**_ to indicate the position-dependent configurational marginal densities of *ρ* and *π*.

### KL divergence as a natural measure of sampling bias

An ideal measure of the discrepancy between the sampled distribution *ρ*_**x**_ and the equilibrium distribution *π*_**x**_ should be “universal” in the sense that driving that measure to zero implies that error in any expectation also goes to zero. It should also be defined for all densities, and not rely on a system-specific choice of observables. One such measure is the Kullback-Leibler (KL) divergence,

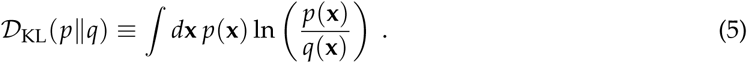

The KL divergence is defined and non-negative for any pair of distributions on the same support, and *𝒟*_KL_(*p*||*q*) = 0 if and only if *p* = *q* almost everywhere.

In [1], Sivak and colleagues demonstrated how to approximate the KL divergence of the sampled distribution *ρ* from the target distribution *π over the full phase-space distribution, 𝒟*_KL_(*ρ*||*π*), in terms of a work-like quantity—the *shadow work*—that is readily computable for a large family of Langevin integrators (Figure 1a). This estimator depends only on the ability to draw samples from *π* and to measure a suitable work-like quantity. This method was applied in [1] to measure the phase-space sampling bias introduced by a particular Langevin integrator (`OVRVO`) on periodic boxes of TIP3P water [22].

**Figure 1.**
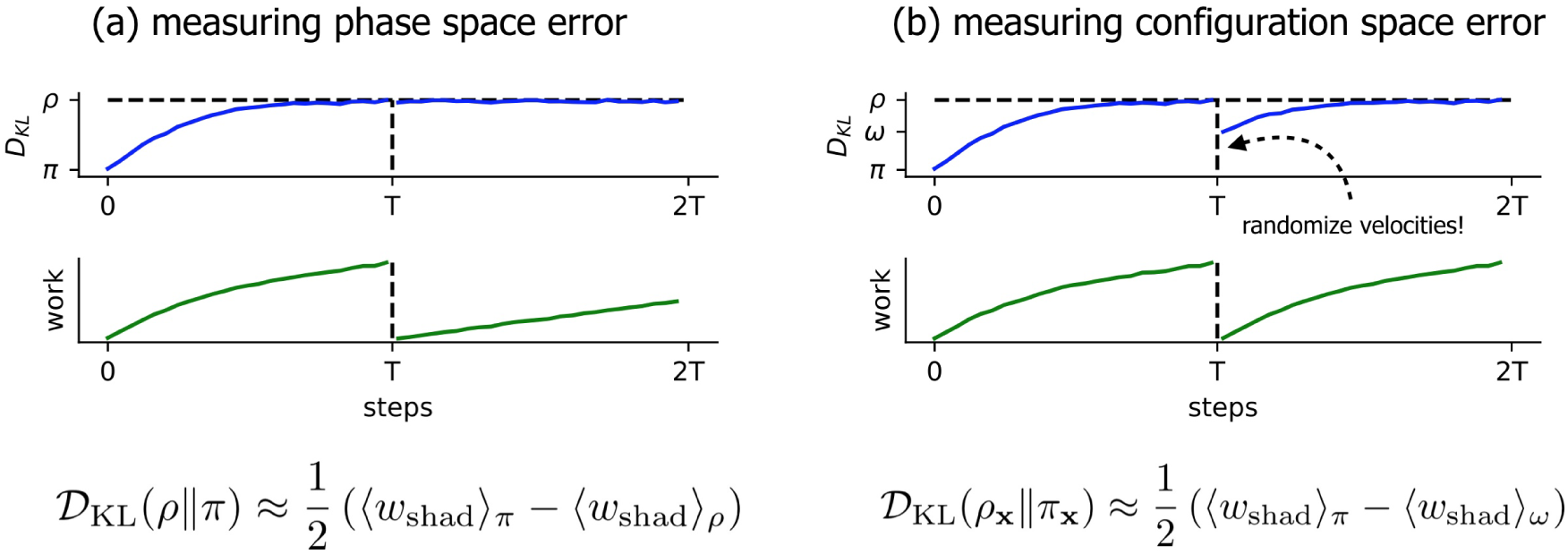
A simple nonequilibrium protocol allows measurement of the KL divergence in phase and configuration space close to equilibrium. Simple nonequilibrium protocols can be used in complex molecular systems to rapidly estimate—utilizing the Crooks fluctuation theorem—the KL divergence of sampled Langevin densities from equilibrium. In both panels, the *x*-axis is the number of steps taken so far in the length-2*T* protocol, and 〈*w*_shad_〉*_π_* indicates the average (reduced, unitless) shadow work accumulated over *T* steps of Langevin dynamics, initialized from equilibrium ((**x**_0_,**v**_0_) ~ *π*). **(a)** The original scheme described in Sivak *et al*. [1] to measure the KL divergence between the sampled phase-space density *ρ* and the equilibrium phase-space density *π*. 〈*w*_shad_)_*ρ*_ is the average shadow work accumulated over *T* steps of Langevin dynamics, initialized from the integrator’s steady state ((**x**_0_,**v**_0_) ~ *ρ*). **(b)** The modified scheme introduced here to measure the KL divergence in the *configuration-space* marginal density between the marginal sampled configuration-space density *ρ*_**x**_ and marginal equilibrium density *π_**x**_*. 〈*w*_shad_〉_ω_ is the average shadow work accumulated over *T* steps of Langevin dynamics, where the initial configuration is drawn from the integrator’s steady state, and the initial velocities are drawn from equilibrium (**x**_0_ ~ *ρ*_x_, **v**_0_ ~ *π*(**v**|**x**_0_)). We denote this distribution *ω*(**x,v**) = *ρ*_x_(**x**)*π*(**v|x**). The **top row** schematically illustrates “distance from equilibrium”, with *y*-axis ticks for *𝒟*_KL_ (*π*||*π*) = 0, *𝒟*_KL_ (*ω*||*π*) ≤ *𝒟*_KL_ (*ρ*||*π*). The **bottom row** illustrates the average work (here, just shadow work) accumulated throughout each protocol.

Since the velocity marginal is not generally of interest, and since some integrators are thought to preserve the configuration marginal of the target distribution with higher fidelity than the phase-space joint distribution, we sought to generalize the technique to estimate the KL divergence of the sampled configuration-space marginal, *𝒟*_KL_ (*ρ*_**x**_||*π*_**x**_). Below, we show how a simple modification of the estimator described in [1] can achieve this goal, and illustrate how this provides a useful tool for measuring the integrator-induced error in configuration-space densities for real molecular systems.

## 2. Numerical discretization methods and timestep-dependent bias

There are several possible ways to discretize Langevin dynamics. A flexible approach to this task is via operator splitting, where the Langevin system is split into components, for example,

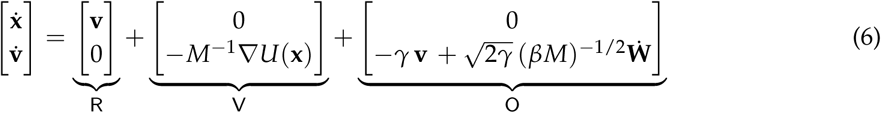

where each component can be solved “exactly” (in the sense of distributions) for a small time increment. The label `O` indicates that the corresponding part of the splitting has the form of an Ornstein-Uhlenbeck process. The labels of the deterministic parts, `R` and `V`, have been chosen to reflect the deterministic updates of position and velocity, respectively; this notation has been used in some previous articles [23, 24]. Note that in the articles of Leimkuhler and Matthews [4,14], the Langevin equations are cast in position-momentum form instead of position-velocity form and the labels `A, B` are then used to indicate the deterministic updates of positions and momenta, respectively. The choices of components of the splitting we use here are not the only options. For example the Stochastic Position Verlet method [25] groups together both of the contributions `V` and `O`. Other splitting-based methods for Langevin dynamics are discussed for example in [26–28].

Once a splitting is defined, the propagator *e* ^ℒΔ*t*^ can be approximated as a Trotter factorization, i.e. a product of the propagators corresponding to the individual components, defining thus a family of numerical methods indexed by the string indicating the order of appearance of these individual terms. For example, we would use `OVRVO` to refer to the composition method,

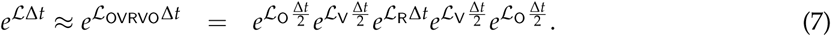

Due to the lack of commutativity of the operators, equality between the true propagator and the Strang splitting is only achieved in the limit Δ*t* → 0, i.e., for vanishing timestep. However, in the case of splitting methods as defined above, it is possible to analyze the error in the effective probability distribution sampled by the finite timestep method [29].

The advantage of the splitting approach is that each component propagator, 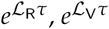, and 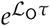, has a straightforward interpretation in terms of arithmetic operations:

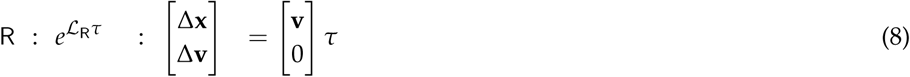

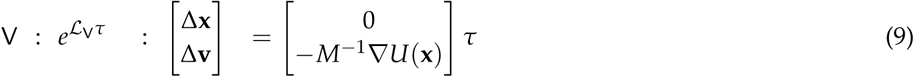

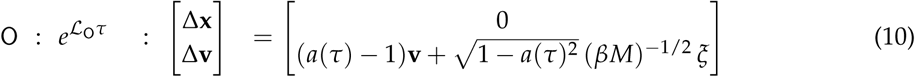

where *a*(*τ*) = *e*^−*γτ*^ and *ξ* ~ 𝒩(0,1)^3*N*^ is a vector of standard normal random variables drawn for each degree of freedom in each `O` step.

By chaining the operations in the order specified by the splitting string, we can unroll them into the sequence of mathematical updates needed to implement one cycle of the integrator for a total time Δ*t*. For `VRORV`, for example, translating the splitting string into the appropriate sequence of update equations in (8)–(10) produces the following equations for one complete integrator timestep:^2^

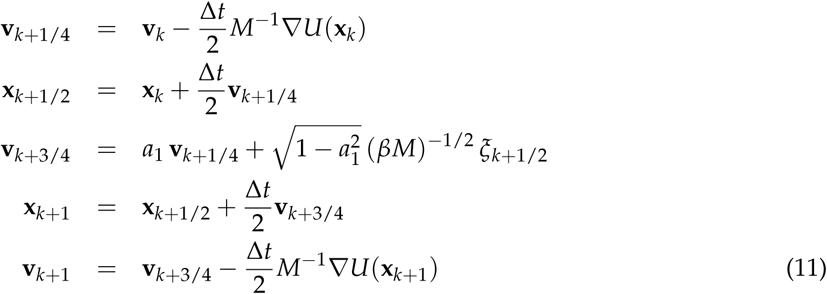

Due to the different naming convention they adopt, this is referred to as the “BAOAB” method in the work of Leimkuhler and Matthews [4,14].

As another example, the Langevin integrator of Bussi and Parrinello [12] corresponds to the splitting `OVRVO`; this splitting is also known as velocity Verlet with velocity randomization [24] due to its use of a velocity Verlet integrator core (substeps `VRV`) [30].

While both the `VRORV` and `OVRVO` discrete time integration schemes reduce to the same stochastic differential equations in the limit that Δ*t* → 0, they can behave quite differently for finite timesteps (Δ*t* > 0), especially for timesteps of practical interest for atomistic molecular simulation.

### Langevin integrators introduce sampling bias that grows with the size of the timestep

In many types of molecular simulations, only configuration-space properties are of interest. The configurational canonical density is defined by marginalization, viz,

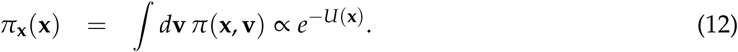

In practice, the velocities are simply discarded in computations while the positions are used to estimate configuration-dependent properties.

The continuous-time Langevin equations of motion (4) possess the target equilibrium density *π* (3) as their unique stationary density, suggesting that, at least in principle, temporal averages along Langevin trajectories can be used to approximate averages with respect to the equilibrium distribution. However, numerical simulations with a finite timestep Δ*t* > 0 will generally sample a different distribution, which we will denote by *ρ*(**x, v**), which implicitly depends on timestep Δ*t*. The discrepancy between the distributions *ρ* and *π* will grow with the size of Δ*t* according to some power law (*e.g*., 𝒪(Δ*t^2^*) or 𝒪(Δ*t^4^*)). For sufficiently large stepsize (typically inversely proportional to the fastest oscillatory mode present) the discretization will become unstable, but the value of the stepsize for which the bias is unacceptable may occur well below the molecular dynamics stability threshold [14].

Note that this phenomenon is completely separate from numerical issues introduced by finite-precision arithmetic on digital computers, which introduces roundoff error in mathematical operations; here, we presume that computations can be carried out to arbitrary precision, and analyze only the effects of time discretization. The timestep-dependent bias is also unrelated to the Monte Carlo or sampling error which is due to finite approximation of the long-term average of a given quantity.

In Figure 2, we illustrate a few key behaviors of this stepsize-dependent sampling bias in a simple quartic 1D system. Note that: (1) the numerically sampled distribution deviates from the target distribution, (2) this deviation increases with timestep Δ*t*, and (3) the deviation in phase space (**x, v**) may be different than the deviation in configuration space only (**x**).

**Figure 2.**
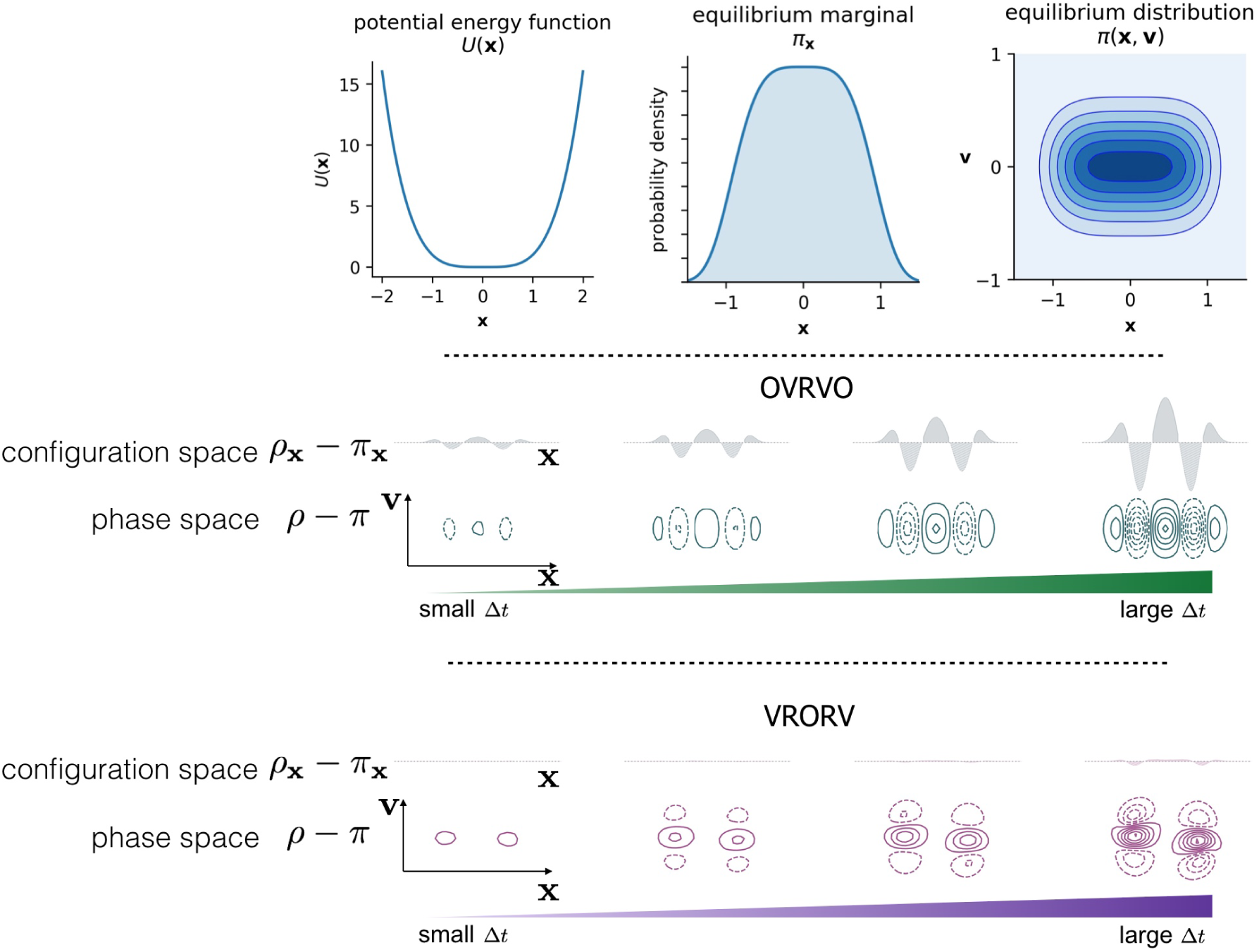
Comparison of Langevin integrators in terms of phase-space and marginal distributions. For a simple 1D system with the quartic potential 𝒰(**x**) = **x**^4^, the error in sampled phase-space density *ρ* and its marginal density *ρ*_**x**_ grows as a function of timestep Δ*t*. However, different Langevin integrators (`OVRVO` and `VRORV` shown here) derived from symmetric Strang splittings can lead to drastically different error structures in phase space, which can induce fortuitous cancellation of error in the marginal distribution under certain circumstances (`VRORV`), see [20]. In the **top row**, we illustrate the definition of the 1D system **(left:** the potential energy function, 𝒰(**x**) = **x**^4^; middle:the equilibrium marginal density over configuration space, *π*_**x**_ (**x**) ∝ *e*^−β𝒰(**x**)^; **right:** the equilibrium joint distribution over phase space *π*(**x, v**)). In the **middle row**, we illustrate the *increasing discrepancy between the sampled distribution ρ and the equilibrium distribution π*, for both the full phase-space and the marginal configuration space, as a function of timestep Δ*t*, for the given model problem and the particular choice of the Bussi-Parinello Langevin integrator `OVRVO` (7). Here the difference between exact and discrete configurational measures are plotted above the contours of the phase space density, for four values of the stepsize Δ*t* = [0.43, 0.66, 0.88,1.1]. In the **bottom row**, we illustrate the timestep-dependent error in a similar way for another integrator `VRORV` (11).

**Figure 3.**
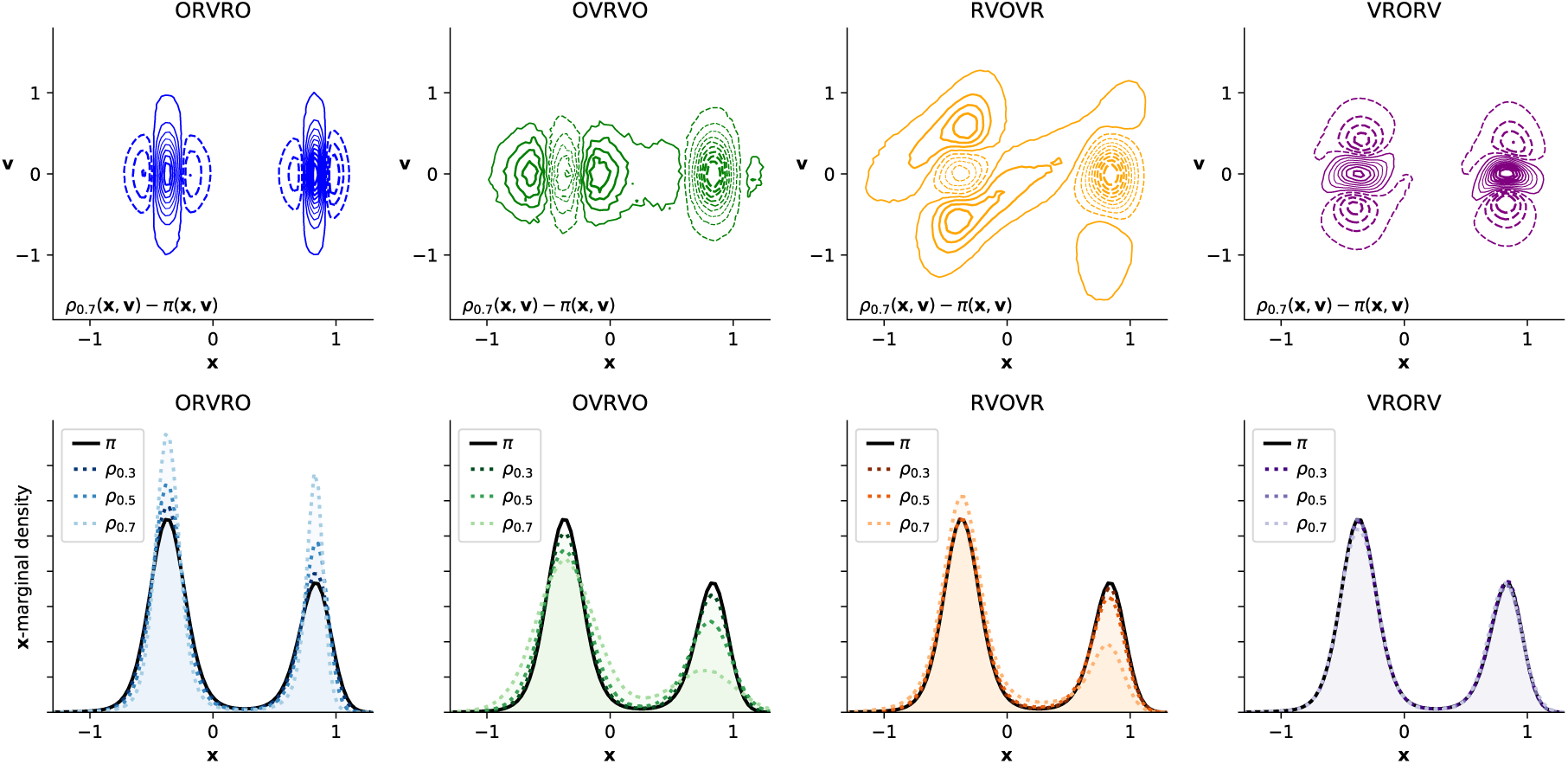
Different numerical integrators introduce different error structure in phase space, illustrated in a double-well system. Here, we illustrate the timestep-dependent discretization error introduced by four integrators on a 1D double-well potential [𝒰(**x**) ≡ **x**^6^ + 2 cos(5(**x** + 1))]. The **top row** of 2D contour plots illustrates the difference between the phase-space density *ρ*(**x, v**) sampled at the maximum timestep considered (Δ*t* = 0.7, close to the stability limit) and the equilibrium density *π*(**x, v**); solid lines indicate positive contours, while dashed lines indicate negative contours. The **bottom row** of 1D density plots shows timestep-dependent perturbation in the sampled marginal distribution in configuration space, *ρ*_**x**_, with the equilibrium distribution *π*_**x**_ depicted as a solid black line. The sampled marginal distributions *ρ*_**x**_ are shown for increasingly large timestep, denoted *ρ*Δ*_t_*, depicted by increasingly light dotted lines, for Δ*t* = 0.3,0.5,0.7 (arbitrary units). Inspecting the contour plots suggests that some integrator splittings (especially `VRORV`) induce error that fortuitously “cancels out” when the density is marginalized by integrating over v, while the error in other integrator splittings (ORVRO, `OVRVO`) constructively sums to amplify the error in configuration space.

It has been proposed that some integrators of Langevin dynamics (particularly the `VRORV` aka “BAOAB” integrator of Leimkuhler and Matthews) preserve the configuration distribution with significantly greater fidelity than other equal-cost integration algorithms [14,31,32], a property that could have significant ramifications for the efficiency of sampling given fixed computational budgets. However, as the formal arguments for this “superconvergence” property rely on a high-friction limit, it is unclear how large the friction coefficient needs to be in practice for the argument to apply. Formal descriptions of the error are typically generic, in that they do not provide guidance on precisely which Δ*t* introduces a tolerable amount of bias for a particular system, and they do not provide a way to predict how other choices, such as mass matrix modifications (*e.g*., hydrogen mass repartitioning) [17,33–35], will affect the error for a system of interest.

## 3. Estimators for KL divergence and the configurational KL divergence

### 3.1. Near-equilibrium estimators for KL divergence

By using work averages, Sivak and Crooks [36] derived a near-equilibrium estimator for the KL divergence between an arbitrary distribution *ρ* and the equilibrium distribution *π*:

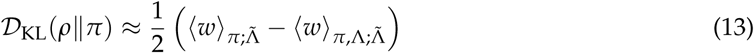

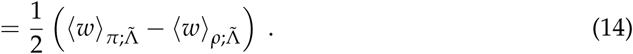

Here 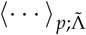 indicates an average over the dynamical ensemble produced by initialization in microstates sampled from density *p* and subsequent driving by protocol 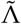 that is the time-reversal of protocol Λ. The expectation 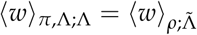 represents a procedure in which initial microstates are sampled from an initial density *ρ* = (*π*,Λ) prepared by sampling *π* and applying protocol Λ; the expectation is subsequently computed by averaging the work during the application of the time-reversed protocol 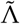 over many realizations of this sampling process. Note that this is distinct from 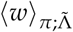, the expectation where the initial sample is selected from *π* and the work is measured during the execution of time-reversed protocol 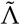.

Sivak, Chodera, and Crooks [1] demonstrated how to apply this estimator when *ρ* is the biased (nonequilibrium) stationary distribution resulting from protocol Λ, the repeated application of a particular numerical Langevin integrator for sufficiently long to reach steady state. In particular, because the numerical Langevin integrator is symmetric (so 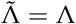) and because the time-independent Hamiltonian produces no explicit protocol work (so *w* = *w*_shad_), in this case the KL divergence is approximately

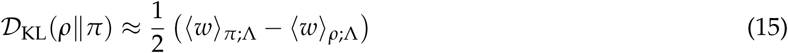

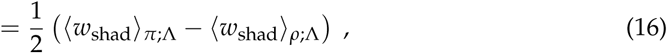

the halved difference of two work averages: the work 〈*w*_shad_〉_*π*_ required to drive from equilibrium *π* into the steady state *ρ*, and the steady-state work 〈 *w_shad_*〉 _*ρ*_ expended over the same length of time, but starting in *ρ* (Figure 5a)^3^.

**Figure 4.**
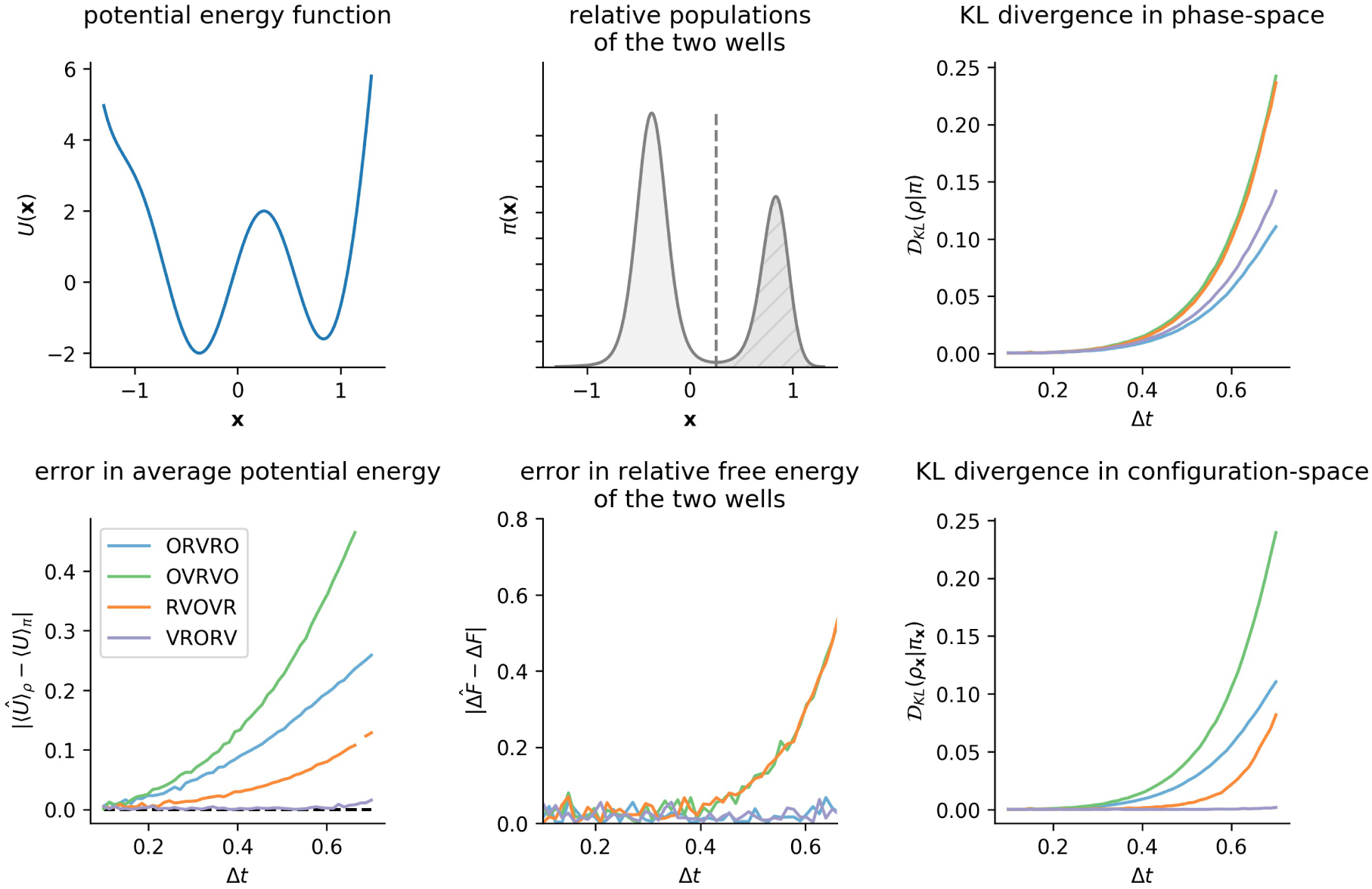
KL divergence is a natural measure of sampling error, although system-specific observables display different sensitivities to sampling error. Even for the simple double-well potential considered in Figure 3, configuration-space properties display different sensitivities to sampling error, motivating the use of a “universal” error measure, such as the KL divergence. The **top left** panel illustrates the double-well potential energy function from Figure 3, and the **top center** panel shows the resulting marginal equilibrium density, *π*_**x**_, at *β* = 1. The **bottom left** panel shows, as a function of Δ*t*, growth in the magnitude of the error in average potential energy, ∣〈𝒰〉*_ρ_* – 〈𝒰〉*_π_*∣, which has been used previously as a sensitive measure of sampling error [14]. The **bottom center** panel shows the error in the apparent free energy difference between the two wells as a function of Δ*t*. Note that the timestep-dependent behavior of these two observables imply different rankings of integrator fidelity that may mislead one into believing error in *all* observables remains low with increasing timestep. However, as is clear here, just because an integrator introduces low timestep-dependent error in one observable does not mean that the method will introduce low error in another observable: for example, `OVRVO` preserves the well populations as accurately as `VRORV`, but introduces much larger errors in the average potential energy. The **right column** summarizes the growth in timestep-dependent error, as measured by the KL divergence. While all four integrators introduce comparable levels of Δ*t*-dependent error in the phase-space distribution, they induce dramatically different magnitudes of error in the configuration-space marginal.

**Figure 5.**
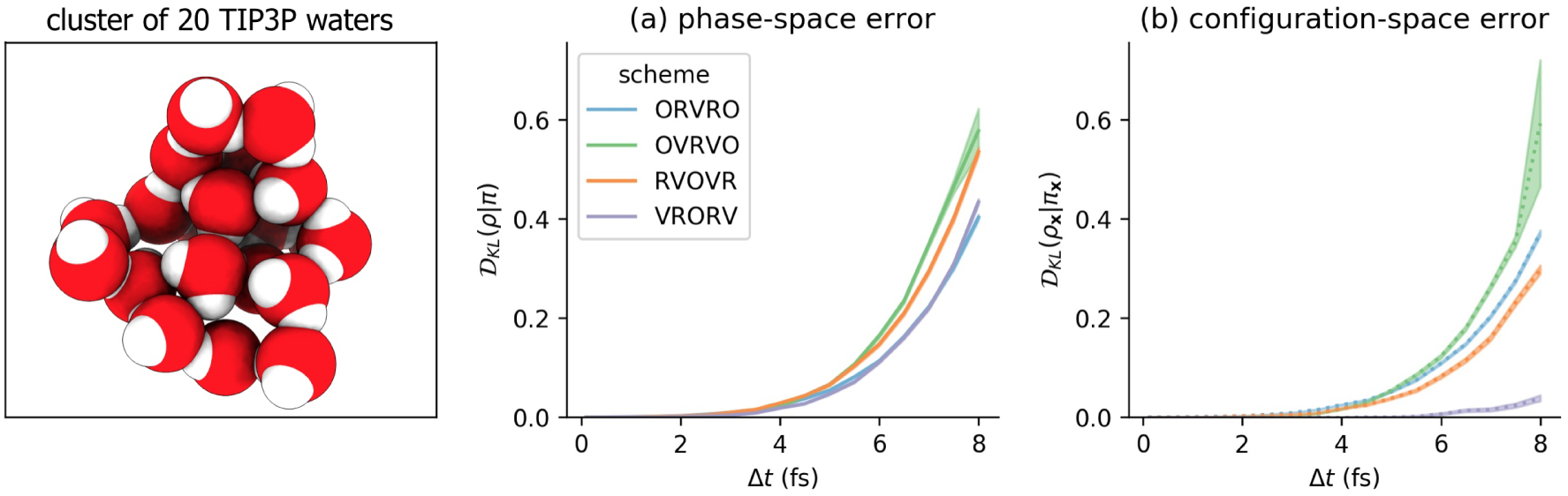
Using the near-equilibrium approximation, some numerical methods introduce far less configuration-space bias in molecular mechanics models than others. The results here are reported for a small cluster of rigid TIP3P waters, described in more detail in the Detailed Methods section, and illustrated **in the leftmost panel**. On the *x*-axis is the timestep Δ*t*, measured in femtoseconds (fs). On the *y*-axis is the estimated KL divergence 𝒟_KL_. (a) The error over the joint distribution on 𝒟_KL_(*ρ*||*π*) (b) The error over the configuration-space marginal 𝒟_KL_(*ρ*_**x**_||*π*_**x**_). Each colored curve corresponds to a numerical scheme for Langevin dynamics. The shaded region is the mean ± 95% confidence interval.

### 3.2. A simple modification to the near-equilibrium estimator can compute KL divergence in configuration space

In this study, we are especially interested in the configuration-space marginal distribution 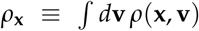. The KL divergence 𝒟_KL_(*ρ*_**x**_||*π*_**x**_) between the respective configuration-space marginal distributions *ρ*_**x**_ and 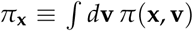 equals 𝒟_KL_(*ω*||*π*), the KL divergence between the full equilibrium distribution *π* and the distribution

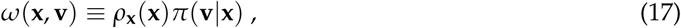

that differs from *π* only in its **x**-marginal:

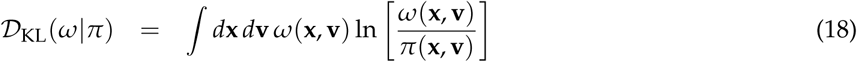

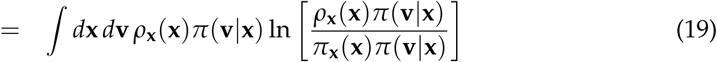

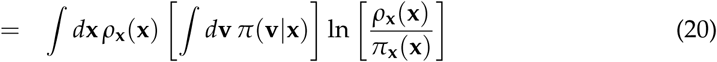

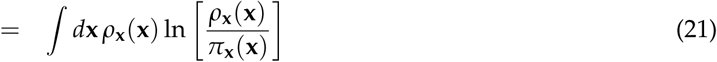

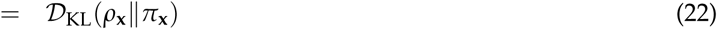

This distribution *ω* is reached via an augmented protocol Λ′ consisting of the original protocol Λ (repeated application of the numerical Langevin integrator) followed by final randomization of velocities according to the equilibrium conditional distribution *π*(**v|x**). Thus we construct an analogous near-equilibrium estimator of 𝒟_KL_(*ω*||*π*) based on average the average shadow work *w*_shad_ accumulated by trajectories initiated from the respective distributions:

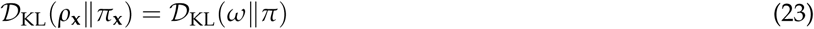

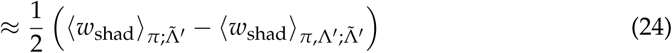

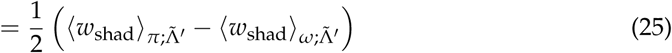

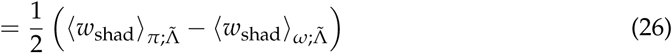

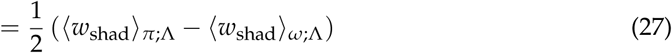

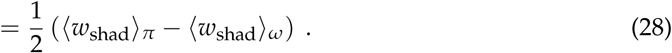

Equation (25) follows from (26) because applying the time-reversed protocol 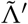 (velocity randomization followed by repeated application of the numerical Langevin integrator) to *π* or to *ω* is equivalent to applying 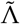: *π* and *ω* already have velocity distributions randomized according to *π*(**v|x**), and the velocity randomization step does no work. In equation (28), we suppress the explicit protocol dependence, since henceforth all work averages will be averaged over the same protocol Λ.

Compared to the full-distribution KL divergence, the configuration-only KL divergence replaces the second work average 〈*w*_shad_〉_*ρ*_ with 〈*w*_shad_〉_*ω*_, computing the expectation of the shadow work over a modified initial density constructed from the nonequilibrium steady-state configuration distribution but with Maxwell-Boltzmann velocity distribution *π*(**v|x**). Practically, this corresponds to drawing samples from the nonequilibrium steady-state *ρ* and replacing the velocities v with an i.i.d. sample from *π*(**v|x**). The modified procedure is depicted schematically in Fig. 1b.

### 3.3. Comparison of phase-space error for different integrators

We first applied the original near-equilibrium method of Sivak *et al*. [1] to measure the timestep-dependent phase-space error introduced by four common Langevin integrators on a molecular mechanics model system (a cluster of 20 TIP3P waters [22] in a harmonic restraining potential), for a range of timesteps Δ*t* between 0.1 and 8.0 femtoseconds (0.1 fs, 0.5 fs, 1.0 fs, …, 7.5 fs, 8.0 fs). As illustrated in Figure 5a, while this approach can resolve statistically significant differences among schemes, none of the four integrator splitting schemes offers a considerable reduction in phase space error. This may be unsurprising, as none of these integrator schemes, when Metropolized, are known to provide a significant reduction in acceptance rates, which depend in some manner on the induced phase-space sampling error (Figure 8a; see Section 4 for more details about the relationship between Metropolized acceptance rates and KL divergence). Consistent with the results of [1], the phase-space error appears to scale approximately as 𝒪(Δ*t^4^*).

### 3.4. Comparison of configurational KL divergence for different integrators

Figure 5b shows that the measured KL divergence between the configuration-space marginals *ρ*_**x**_ and *π*_**x**_ can be drastically different among the four integrator schemes, and in some cases grow much more slowly than the associated phase-space sampling error (Fig. 5a). In particular, for `VRORV`, the error in the **x**-marginal is very nearly zero for the entire range of feasible timesteps, and it can be run at Δ*t* ≈ 6 fs while introducing the same amount of configuration error as other methods at Δ*t* ≈ 2 fs. This is consistent with prior findings [4,14], which showed the `VRORV` scheme introduces very little error in the average potential energy and multiple other system-specific observables.

We also note that for the `OVRVO` scheme, 𝒟_KL_(*ρ*_**x**_||*π*_**x**_) ≈ 𝒟_KL_(*ρ*||*π*) over the range of measured timesteps (Fig. 5), consistent with prior findings that estimates of 𝒟_KL_(*ρ*||*π*) tracked well with several measures of configuration-sampling error [1].

### 3.5. Influence of the collision rate

The results reported above are for a single, relatively weak collision rate of 1 ps^−1^. This collision rate was selected, following prior work [14], because it is low enough that barrier-crossing and conformational exploration should be fast, but large enough that low errors are observed empirically for configuration-space averages. As the formal “superconvergence” properties of `VRORV` were derived in the high-friction limit [14,31,32], it is worth considering how robustly the splitting `VRORV` introduces low configuration-space error at various collision rates. Additionally, as the collision rate goes to zero, We would expect the differences between pairs of the schemes that are equivalent upon removal of the `O` step (such as `OVRVO` and `VRORV`) to become smaller as the collision-rate goes to zero.

In Figure 6, we report near-equilibrium estimates of 𝒟_KL_ (as in Figure 5) over a range of collision rates spanning *γ* from 0.1–100 ps^−1^. Strikingly, the configuration-space error introduced by `VRORV` remains low over this entire range, while other integrators (such as `OVRVO`) display a significant sensitivity to collision rate. In general, increasing collision rate increases phase- and configuration-space error for all integrators for which differences can statistically be differentiated, though the effect is modest over several orders of magnitude relative to the sensitivity with respect to timestep. For each condition, 50000 protocol samples were collected (one half the number of samples used in Figure 5), and the protocol length was 2000 steps (twice the protocol length used in Figure 5). Figure A7 demonstrates that this result is robust to protocol length for all collision rates considered.

**Figure 6.**
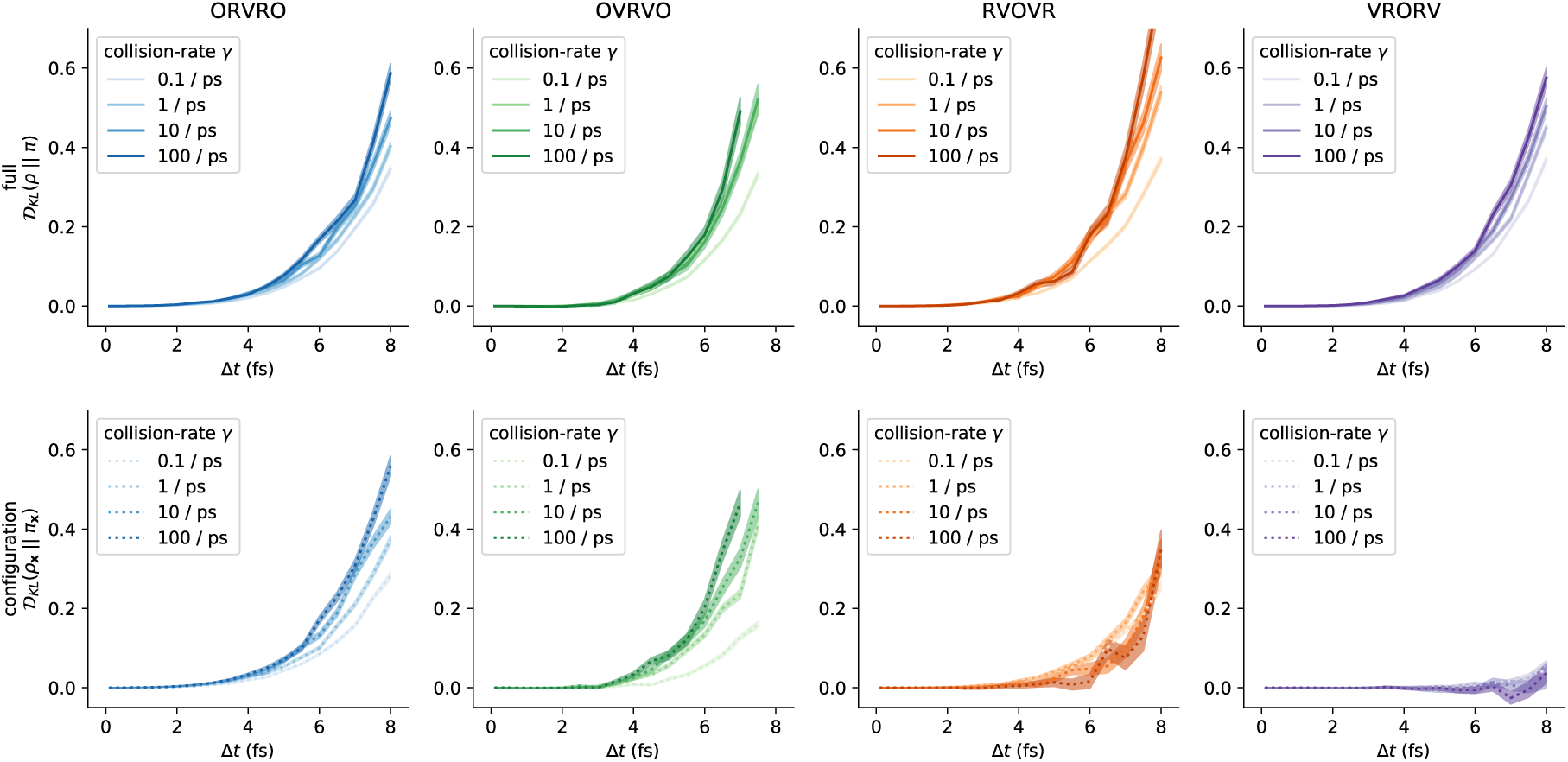
The choice of collision rate influences sampling bias. As we vary the collision rate *γ* over a few orders of magnitude, the resulting measured KL divergence responds in different ways for the different schemes. The phase-space bias appears to increase with increasing collision rate for all schemes. The configuration-space bias for `OVRVO` and `ORVRO` appears to *increase* with increasing collision rate, but the configuration-space bias for `RVOVR` appears to *decrease* with increasing collision rate. The anomalous low configuration-space error for `VRORV` is observed across all collision rates tested. The non-monotonic curves in the *γ* = 100 ps^−1^ condition are expected to be due to finite-sampling error, and are expected to be attenuated at a larger number of protocol samples. (Note that one condition is omitted from these plots for clarity: estimates of 𝒟_KL_ for `OVRVO` at Δ*t* = 8 fs. At that timestep, the variance of the resulting 𝒟_KL_ estimates for this scheme were much larger than for the other schemes.) See Figure A6 for a comparison grouped by collision rate, rather than by integrator.

### 3.6. Comparison with reference methods validates the near-equilibrium estimate

The accuracy of the near-equilibrium approximation introduced by Sivak *et al*. is largely unexplored. While the near-equilibrium approximation is computationally and statistically appealing, it is important to validate the accuracy of the approximation over the practical timestep Δ*t* range of relevance to molecular simulation. In particular, it is unknown whether the near-equilibrium approximation produces an over-estimate or under-estimate of the KL divergence, or how accurate the approximation is for high-dimensional systems. Further, it is unknown whether any bias introduced by the approximation is uniform across different numerical methods for Langevin dynamics.

How well does the near-equilibrium estimator approximate the true KL divergence of relevant timestep ranges? The task of validating the near-equilibrium approximation is numerically challenging, since we are unaware of exact estimators for 𝒟_KL_ (*ρ*||*π*) that remain tractable in high dimensions^4^. In the case of simple fluids, approximate methods are available that express the KL divergence in terms of a series of N-body correlations (as in [38]), typically truncating to two-body correlation functions (*i.e*., comparing the radial distribution functions). However, in general we do not know the effect of truncating the expansion, since the successive terms in the series do not necessarily have decreasing magnitude.

To validate the near-equilibrium estimate of the KL-divergence, we attempt to “sandwich” it between two reference estimates that are unlikely to substantially over- or under-estimate the KL-divergence, and verify that the proposed method is consistent. In Section 3.6.1, we derive an asymptotically exact nested Monte Carlo method, that is computationally inefficient and an under-estimate in practice. In Section 3.6.2, we note that the results from the nested Monte Carlo method can be reprocessed to yield an over-estimate. In Section 3.6.3, we compute both, and compare with the near-equilibrium estimator.

#### 3.6.1. Practical lower bound from nested Monte Carlo

First, we will derive an exact expression for the KL divergence between *ρ* and *π* in terms of quantities that we can measure, then discuss practical challenges that arise when using this expression, and under what conditions it becomes impractical. We start by writing the KL divergence as an expectation over *ρ* (30), since we cannot evaluate ρ(**x,v**) pointwise, but we can draw samples (**x,v**) ~ *ρ*.

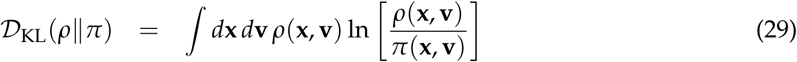

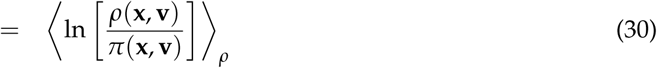

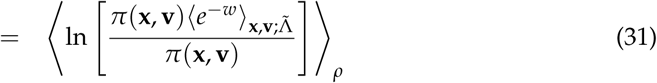

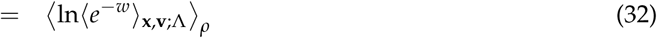

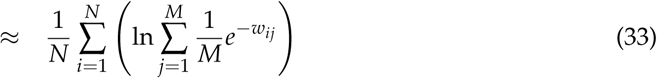

We note that the inner ratio of nonequilibrium steady-state to equilibrium densities, *ρ*(**x,v**)/*π*(**x,v**), can be expressed in terms of 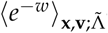—the average of exponentiated nonequilibrium work measured under the application of the time-reversed protocol 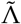 starting from (**x, v**) (31). Λ denotes the protocol used to generate *ρ* from a sample from *π*—in this case, T applications of the Langevin integrator step kernel; 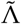 denotes the time-reverse of this protocol. Since the protocol we apply to generate *ρ* from *π* is time-symmetric for integrators derived from symmetric Strang splittings^5^, we can substitute 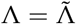. In the final step, we substitute a simple Monte Carlo estimator of that average, in terms of work samples *w_ij_*, where *w_ij_* is the *j*th reduced (unitless) work measurement collected from initial condition *i*. Here, *N* is the number of initial conditions sampled (*i.e*., the number of “outer-loop” samples), and *M* is the number of work samples (*i.e*., the number of “inner-loop” samples) collected at each initial condition (**x***_i_*,**v***_i_*) ~ *ρ*.

Related nested estimators have been proposed in the literature. Notably, an estimator for the nonequilibrium entropy 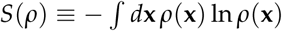 in terms of work averages is given in equation 18 of [39], and this can in turn be used to estimate the desired KL divergence if we also have suitable estimates for the equilibrium entropy *S*(*π*).

The required work values *w_ij_* can be easily computed from simulations. To sample an initial condition (**x***_i_*, **v***_i_*) from *ρ*, we simply run the Langevin integrator of interest for a sufficient number of steps to sample a new uncorrelated configuration from the nonequilibrium steady-state sampled by the integrator. To compute the work accumulated from a given starting condition, we use the notion of *shadow work* [23]. For numerical methods constructed from symmetric Strang splittings involving the `R, V`, and `O` operations described above, we simply need to compute the sum of the total energy changes during the deterministic substeps (*i.e*., the potential energy change during deterministic updates of the position variables, and the kinetic energy change during deterministic updates of the momentum variables). For convenience, we use reduced (unitless) energies and work values throughout, where factors of *k_B_*T have been removed, without loss of generality. See Detailed Methods (Section 6.4) for a detailed description on how shadow work can be computed from this family of Langevin integrators in general.

Like the near-equilibrium scheme, this nested scheme can be modified analogously to measure the configuration-space error in isolation, by initializing instead from the distribution *ω*(**x,v**) ≡ *ρ*_**x**_(**x**) *π*(**v|x**), allowing us to compute 𝒟_KL_(*ω*||*π*), a quantity that is identical to 𝒟_KL_(*ρ*_**x**_||*π*_**x**_) (see 3.2). Specifically, to measure the full KL divergence, we sample initial conditions from the Langevin integrator’s steady state: (**x***_i_*,**v***_i_*) ~ *ρ*. To measure configuration-space-only KL divergence, we draw initial configuration from the integrator’s steady state, and velocities from equilibrium: **x***_i_* ~ *ρ*_**x**_, **v***_i_* ~ *π*(**v**_*i*_|**x**_*i*_). Note that, for constrained systems, *π*(**v|x**) is not independent of **x**, and care must be taken to eliminate velocity components along constrained degrees of freedom before measuring the contribution of the integrator substep to the shadow work (see Detailed Methods).

We note that the nested plug-in Monte Carlo estimator of the KL divergence is asymptotically exact only when both *N* (the number of “outer-loop” samples) and M (the number of “inner-loop” samples) go to infinity. For a finite number of samples, the nested estimator will produce an under-estimate of the KL divergence. To see this, note that the nested estimator is a simple average of many likely underestimates, so it should itself be an underestimate. More specifically, although the outer-loop expectation can be approximated without bias (since samples can be drawn from *ρ*), each of the inner-loop expectations is an exponential average (log〈exp(−*w*)〉), which we will underestimate when we plug in a finite sample of *w’s* (for the same reasons that the EXP estimator for free energies is biased [40] – To leading order, that under-estimate is related to the variance of *w* [40], which here grows rapidly with Δ*t* as shown in Figure A2).

In practice, we use a simple adaptive scheme (described in detail in Section 6.6) that draws inner-and outer-loop samples until uncertainty thresholds are met, which should minimize the magnitude of this bias.

#### 3.6.2. Practical upper bound from Jensen’s inequality

A simple, but practically useful, upper bound for the KL divergence can be obtained from the application of Jensen’s inequality, 〈ln *x*〉 ≤ ln 〈*x*〉, to (32):

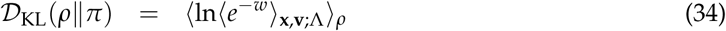

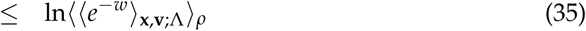

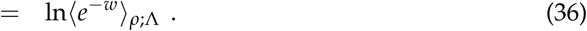

The analogous inequality for the configuration-space marginal is

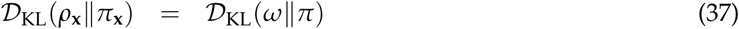

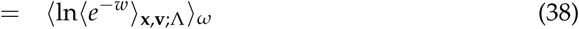

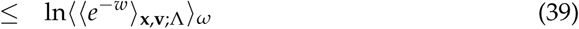

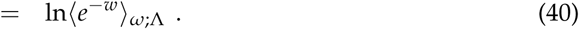

This provides a particularly convenient upper bound on the KL-divergence, since it can be computed by reprocessing work samples collected from the nested Monte Carlo scheme^6^.

#### 3.6.3. Sandwiching the KL divergence to validate the near-equilibrium estimate

We compared these three estimates of the KL divergence on the molecular mechanics system introduced in Figure 5, and confirmed that the near equilibrium estimate falls between the likely over-and under-estimate for all four integrator schemes, over a range of feasible timesteps (Figure 7, and in log-scale in Figure A5). We conclude that the near-equilibrium approximation is empirically reliable for measuring integrator bias on molecular mechanics models for practical timesteps.

**Figure 7.**
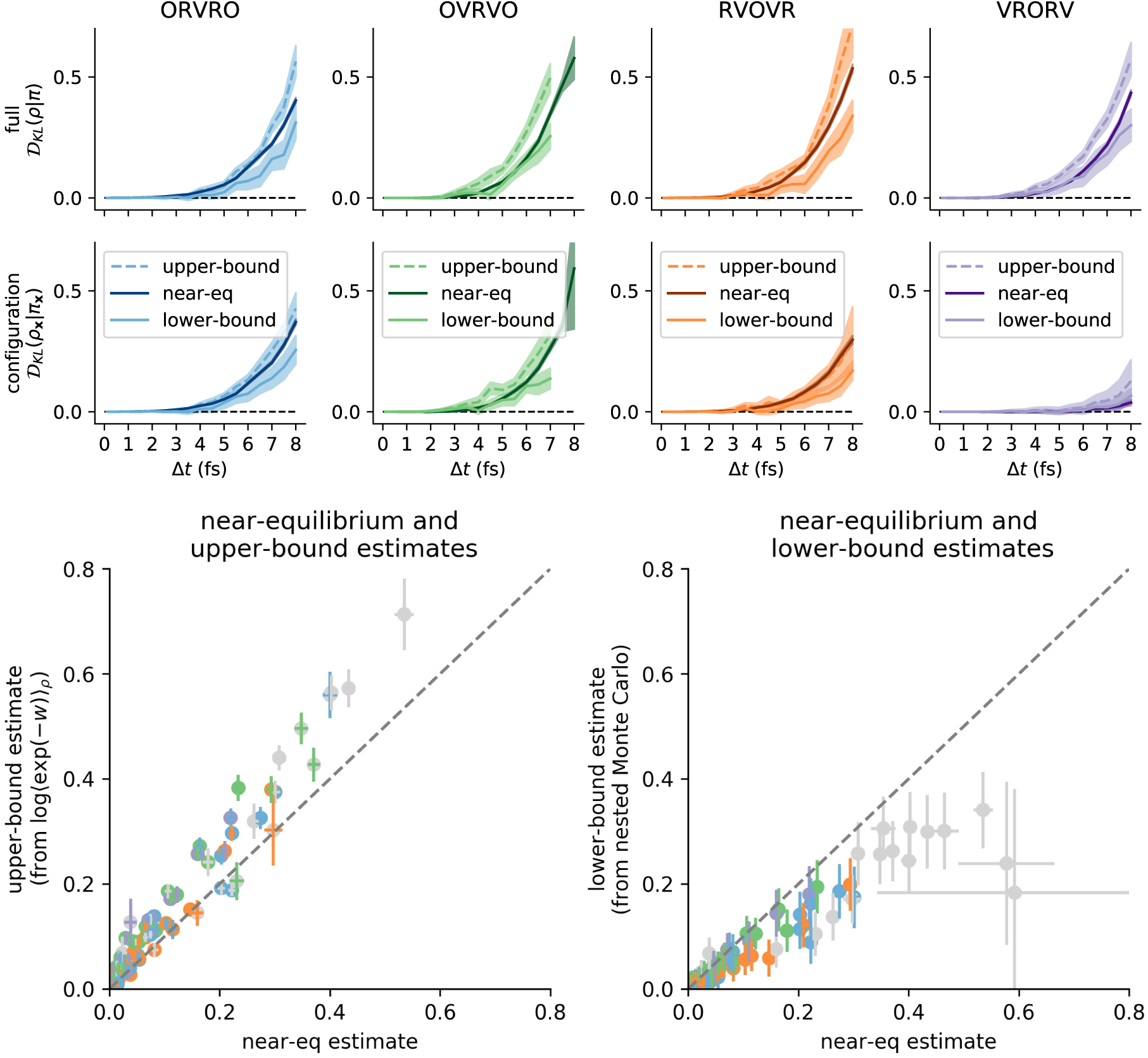
The near-equilibrium estimator is consistent with reference estimators for a practical range of Δ*t*. We compared the near-equilibrium estimates reported in Figure 5 for the water cluster against a likely under-estimate and a likely over-estimate of the 𝒟_KL_. In the **top row**, we validate near-equilibrium estimates of the KL divergence on the full state space (**x**, **v**). In the **bottom row**, we validate near-equilibrium estimates of the KL divergence on configuration space (**x**) alone. Each column corresponds to a numerical method for Langevin dynamics. The **darker band** in each plot corresponds to the near-equilibrium estimate ± 95% confidence intervals from asymptotic uncertainty estimate (details in section 3.1). The **lighter band with a solid line** corresponds to the nested Monte Carlo estimate ± 95% confidence intervals from bootstrapping (details in section 6.6). The **lighter band with a dotted line** corresponds to the exponential average estimate ± 95% confidence intervals from bootstrapping (details in section 6.6). Log-scale versions of these plots are provided in the appendix also, A5. In the **lower two panels**, we summarize these results by plotting all near-equilibrium estimates vs. all exponential-average estimates (left) and all near-equilibrium estimates vs. all nested Monte Carlo estimates (right). The colored dots and bars correspond to the means ± uncertainties used in the earlier panels. The dashed diagonal line shows parity. Grey error dots and error bars correspond to conditions where the nested Monte Carlo estimate reached the computational budget (5 × 10^4^ inner-loop samples) but failed to reach the inner-loop uncertainty threshold, and is thus more biased. See Section 6.6 for additional details.

## 4. Relation to GHMC acceptance rates

What is the relationship between the bias introduced by an integrator at steady state, and the acceptance rate of the corresponding Metropolized integrator? Specifically, why not construct a Metropolized version of `VRORV` to guarantee samples are drawn appropriately from the equilibrium target density *π*(**x**)? Following [3,41], we can construct an exact MCMC method that uses one or more steps of Langevin dynamics as a proposal, by using

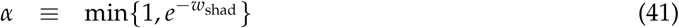

as the acceptance criterion. The resulting method is called *generalized hybrid Monte Carlo* (GHMC), and eliminates time-discretization error at the cost of increasing sample autocorrelation (see also the comments in Section 6.3). A natural question arises: if an uncorrected Langevin integrator introduces low configuration-space error, is the rejection rate of the corresponding Metropolis-corrected method also low?

To answer this question, we estimated the GHMC acceptance rate at all conditions for which we have estimated steady-state 𝒟_KL_. Given a collection of equilibrium samples (described in Section 6.3), we can efficiently estimate the acceptance rate of an MCMC proposal by taking the sample average of the acceptance ratio *α* over proposals originating from equilibrium, (**x**_0_, **v**_0_) ~ *π*.

We compared the GHMC acceptance rate to the histogram-based 𝒟_KL_ estimates for the 1D double-well system in Figure 8. There does not appear to be a consistent relationship between 𝒟_KL_ and acceptance rate across the four schemes. Notably, the GHMC rejection rate can be extremely “conservative” for splittings such as `VRORV`.

**Figure 8.**
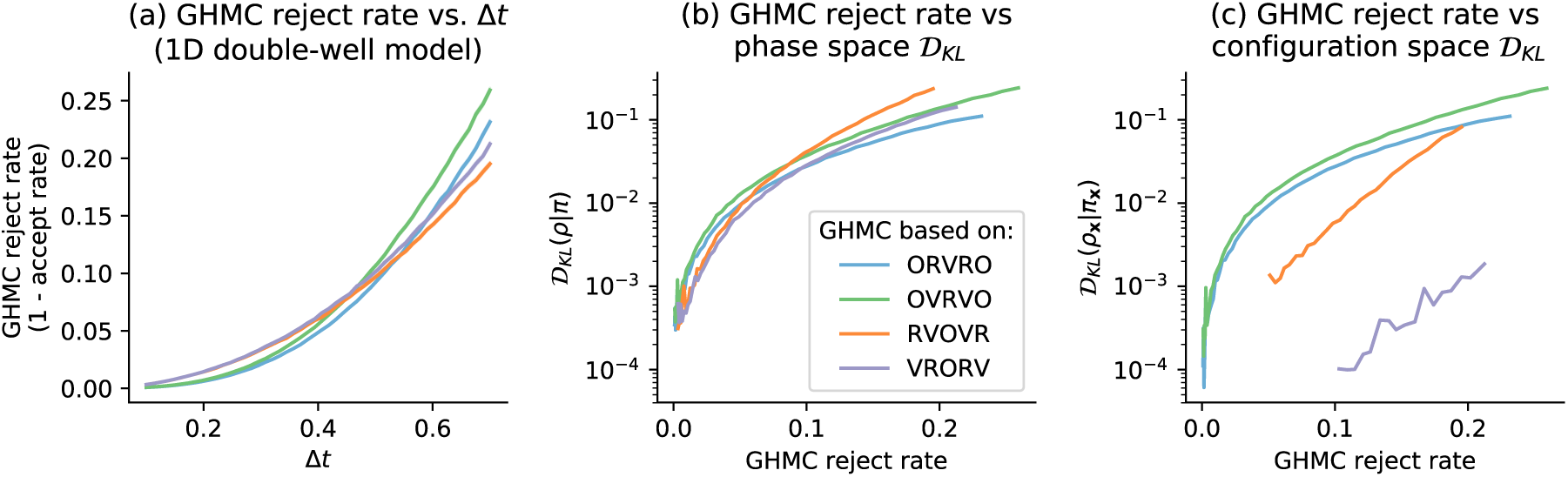
No consistent relationship between the GHMC acceptance rate and the steady-state bias is apparent for a 1D system. Since the GHMC rejection rate grows similarly with Δ*t* across all four schemes, but the configuration-space KL divergence does not, the GHMC rejection rate can be overly “conservative” for some splittings. Panel **(a)** shows the growth in the GHMC rejection rate as a function of timestep Δ*t*, for the 1D double-well model considered in Figures 3 and 4. On the *x*-axis is an evenly spaced grid of 50 timesteps between 0.1 and 0.7. On the *y*-axis is the estimated rejection rate, which is based on a sample average of the GHMC acceptance criterion. The shaded region is the mean ± 95% confidence interval. Panel **(b)** compares the GHMC rejection rate vs. the *phase-space* bias at steady state, over the range of timesteps plotted in panel (a). The *y*-axis is KL divergence between the phase-space histograms, plotted on a log-scale. Panel **(c)** compares the GHMC rejection rate vs. the *configuration-space* bias at steady state, over the range of timesteps plotted in panels (a), (b). The *y*-axis is the KL divergence between the configuration-space histograms, plotted on a log-scale. Note that in panel (c), we have truncated the leftmost parts of the curves for `RVOVR` and `VRORV` rejection rates less than 0.05 and 0.1, respectively, due to noise in histogram estimates of very small 𝒟_KL_(*ρ*_**x**_||*π*_**x**_).

Next, we compared the GHMC rejection rate with the near-equilibrium 𝒟_KL_ estimates for the water cluster considered in Figure 9. A similar pattern is recapitulated in this molecular mechanics model as in the 1D system—there is not a consistent relationship between configuration-space bias introduced by a Langevin integrator and the rejection rate of its corresponding GHMC method.

**Figure 9.**
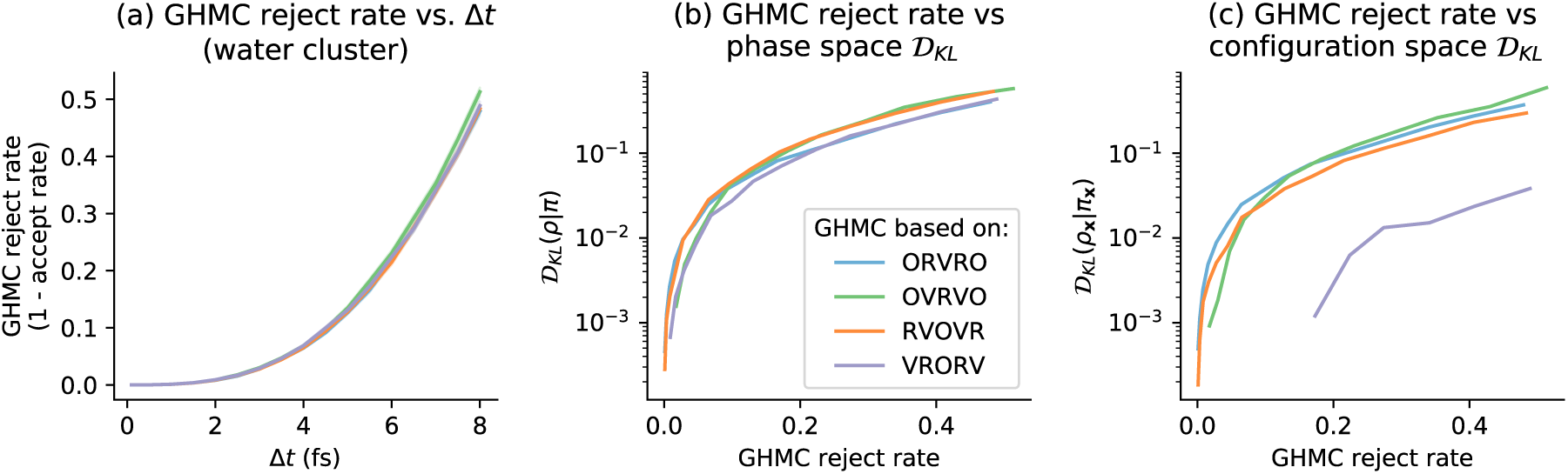
Near-equilibrium measurements recapitulate the relationship between steady-state 𝒟_KL_ and GHMC acceptance rate for the water cluster test system. Panel **(a)** shows the growth in the GHMC rejection rate (1 minus the acceptance rate) as a function of timestep Δ*t* (in femtoseconds), for the water cluster test system illustrated in Figure 5. On the *x*-axis are timesteps [0.1 fs, 0.5 fs, 1.0 fs, … 7.5 fs, 8fs]. On the *y*-axis is the estimated rejection rate, which is based on a sample average of the GHMC acceptance criterion, over 10000 proposals per condition. The shaded region is the mean ± 95% confidence interval. Panel **(b)** compares the GHMC rejection rate vs. the *phase-space* bias at steady state, over the range of timesteps plotted in panel (a). The *y*-axis is the KL divergence between the phase-space distributions as measured by the near-equilibrium estimate, plotted on a log-scale. Panel **(c)** compares the GHMC rejection rate vs. the *configuration-space* bias at steady state, over the range of timesteps plotted in panels (a), (b). Note that in panels (b) and (c), we have truncated at 𝒟_KL_ ≤ 10^−4^, due to noise in near-equilibrium estimates of very small 𝒟_KL_.

This complicates the decision of whether to Metropolize or not. As noted in Section 6.3, incurring even a small rejection rate in GHMC can have a large effect on statistical efficiency, due to the effect of momentum flipping. An open challenge is to construct Metropolis criteria for GHMC that might be less “wasteful” for Langevin splittings that introduce low configuration-space bias. One possibility is to use a multi-proposal Metropolization procedure to correct estimates of equilibrium expectations, as done in [42].

## 5. Discussion

We have introduced and validated a work-based estimator of the KL divergence over the configuration-space marginal sampled by Langevin dynamics. We demonstrated that we could use this estimator to measure differences between the timestep-dependent configuration-sampling error introduced by each member of a small class of Langevin integrators on molecular mechanics models. Work-based estimators are especially attractive for biomolecular systems, since expectations over work distributions are often tractable when other approaches are not.

Reliable estimates of KL divergence using the work-based estimator considered here require knowledge of the time to reach nonequilibrium steady state. This near-equilibrium approach requires that the user select a trajectory length *T* sufficiently large to reach the nonequilibrium steady state, or else the KL divergence estimate could be substantially biased. Opposing the choice of large *T* is the variance of the estimate, since the contribution of the steady-state work to the variance of the estimate grows as *T*. Taken together, this suggests the smallest time *T* that produces unbiased estimates is optimal. In our calculations, it was sufficient to use a protocol that was twice the average collision time, but in general this choice should be validated for the system under study. One way to do this, for example, is to perform the same computation for *T* and 2*T* and ensure estimates are concordant to within statistical error.

Generating equilibrium samples from *π* (**x**, **v**) can be difficult for large, complex molecular systems. The near-equilibrium method requires access to a large number of independent samples from the equilibrium distribution of the system under study. In this work, we used extra-chance HMC [41,43] to construct a large cache of independent equilibrium samples, amortizing the cost of equilibrium sampling across the many integrator variants (described in Section 6.3). In cases where we would like to compare a large number of integrator variants on the same system, this can be an acceptable cost, but in other cases, it may be prohibitive. It is unknown whether we can relax this requirement in practice, and use only samples in “local equilibrium,” since integrator error may be dominated by local features of the energy landscape. An alternative would be that, if the primary contributions to the dissipation process that drive integration errors arise from high-frequency motions, adding a weak restraint to parts of the system under study (such as a biological macromolecule) may also allow rapid assessment of integrator schemes in a region of configuration space where estimates can be easily converged. We have not yet tested this hypothesis, but if it is possible to relax the requirement of i.i.d. equilibrium samples and retain accurate estimates, then the method will be much cheaper to apply in difficult settings.

### 5.1. Future directions

The validation of the near-equilibrium estimate makes it possible to apply the technique to a systematic comparison of sampling bias in integrator variants and biomolecular systems. Although we considered only four Langevin integrators here, this approach can be applied to any stochastic integrator for which the relative path action can be computed (see [44] for examples of how to compute the relative action for stochastic integrators not necessarily derived from operator splitting).

Independently, the work-based estimate for ln[*ρ***_x_**(**x**)/*π*(**x**)] we used in the expensive lower bound (Section 3.6) could be useful for other analyses. For example, an estimate of ln[*ρ***_x_**(**x**)/*π*(**x**)] could be used to interpret what features of **x** are most distorted by integrator bias, *e.g*., by checking which features of **x** are most predictive of extreme values of ln[*ρ***_x_**(**x**)/*π*(**x**)].

We also note that nothing about the derivation is specific to the partition between configuration degrees of freedom and velocities. We could also use this method to measure the KL divergence over any subset *S* of the state variables **z** = (**x**, **v**), provided we can sample from the conditional distribution for the complementary subset *S*′ of the state variables: *π*(**z**_*S*′_|**z***_S_*). To measure KL divergence over the configuration variables, we need only sample from the conditional distribution of velocities given positions, which is typically tractable. Provided that the required conditional distribution is tractable, this method could also prove useful in contexts other than measuring integrator error.

Finally, in this study we have only considered sampling the canonical ensemble (NVT; constant temperature, particle number, and *volume*), but the isothermal-isobaric ensemble (N*p*T; constant temperature, particle number, and *pressure*) also has wide practical relevance. In OpenMM, the isothermal-isobaric ensemble is simulated by alternating between sampling the canonical ensemble (using thermostatted dynamics, such as Langevin dynamics), and periodically sampling the volume using a molecular-scaling Monte Carlo barostat. For sufficiently infrequent barostat proposals, the bias introduced by interaction between the finite-timestep Langevin integrator and the volume perturbations is expected to be minimal, so applying the proposed near-equilibrium method is expected to approximate well the overall sampling bias^7^.

## 6. Detailed methods

All code used in this paper, along with a manifest of all conda-installable prerequsites and version numbers needed to run the code, is available at https://github.com/choderalab/integrator-benchmark under the permissive Open Source Initiative approved MIT license.

A byproduct of this work is a flexible implementation of Langevin integrators derived from operator splitting for the GPU-accelerated OpenMM molecular simulation framework [45], also available under the MIT license in the openmmtools library: https://github.com/choderalab/openmmtools. This implementation allows the user to specify a Langevin integrator using a splitting string (like `OVRVO`) and can automatically compute shadow work for each splitting.

### 6.1. One-dimensional model system: Double well

For illustration and to have a model system where the exact 𝒟_KL_ was readily computable using histograms, we constructed and analyzed a double-well model in 1D. The potential energy function of this model is 𝒰(*x*) ≡ *x*^6^ + 2 cos(5(*x* + 1)), illustrated in Figure 4. We implemented the four Langevin schemes under study using Numba 0.35.0 [46] for use with 1D toy models. We used a temperature of *β* = 1, a collision rate of *γ*= 10, and a mass of *m* = 10. For these conditions, we found a maximum stable timestep of approximately Δ*t* = 0.7. Histogram-based estimates of the configuration-space density and phase-space density used 100 bins per dimension, where the bin edges were set by bounding box of a trial run at the maximum Δ*t*. The equilibrium density of each bin was computed using numerical quadrature (using the trapezoidal rule, `numpy.trapz`). The KL divergence between a given *ρ* and *π* was then computed using `scipy.stats.entropy` on the histogram representation.

### 6.2. Model molecular mechanics system: A harmonically restrained water cluster

As noted, the exact Monte Carlo method involves exponential work averages, resulting in a statistical inefficiency that grows rapidly with both the size of the system and the distance from equilibrium. Since we are interested in identifying whether the near-equilibrium approximation breaks down over the timestep Δ*t* range of interest to molecular simulations, it is important to be able to compute a reliable estimate of 𝒟_KL_ far from equilibrium. Thus, we aim to select the smallest system we think will be representative of the geometry of molecular mechanics models generally in order to allow the exact estimate to be computable with reasonable computing resources.

To compare the proposed method with a reference estimator, we needed to select a test system which met the following criteria:

1. The test system must have **interactions typical of solvated molecular mechanics models**, so that we would have some justification for generalizing from the results. This rules out 1D systems, for example, and prompted us to search for systems that were not alanine dipeptide in vacuum.
2. The test system must have **sufficiently few degrees of freedom that the nested Monte Carlo estimator remains feasible**. Because the nested estimator requires converging many exponential averages, the cost of achieving a fixed level of precision grows dramatically with the standard deviation of the steady-state shadow work distribution. The width of this distribution is extensive in system size. Empirically, this ruled out using the first water box we had tried (with approximately 500 rigid TIP3P waters [22], with 3000 degrees of freedom). Practically, there was also a limit to how small it is possible to make a water system with periodic boundary conditions in OpenMM (about 100 waters, or 600 degrees of freedom), which was also infeasible.
3. The test system must have **enough disordered degrees of freedom that the behavior of work averages is typical of larger systems**. This was motivated by our observation that it was paradoxically much easier to converge estimates for large disordered systems than it was to converge estimates for the 1D toy system.

To construct a test system that met all of those criteria, we used a `WaterCluster test` system, which comprises 20 rigid TIP3P waters weakly confined in a central harmonic restraining potential with force constant *K* = 1 kJ/mol/nm^2^ applied to all atoms. This test system is available in version 0.14.0 of the `openmmtools` package [47]. Simulations were performed in double-precision using the `Reference` platform in OpenMM 7.2 [48] to minimize the potential for introducing significant round-off error due to finite floating point precision.

### 6.3. Caching equilibrium samples

To enable this study, we attempted to amortize the cost of collecting i.i.d. samples from each test system’s equilibrium distribution *π* and various integrator-and-Δ*t*-specific distributions *ρ*. Since there are many different distributions *ρ*, and all are relatively small perturbations of *π*, we invest initial effort into sampling *π* exhaustively, and then we draw samples from each integrator-specific *ρ* by running the integrator of interest from initial conditions (**x**_0_, **v**_0_) ~ *π*.

For each test system, we pre-computed a large collection of *K* = 1000 equilibrium samples

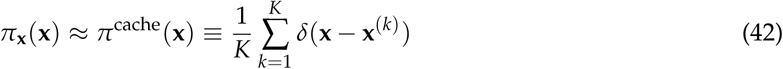

using *extra-chance Hamiltonian Monte Carlo* (XC-HMC)[41,43], implemented as a CustomIntegrator in OpenMM [48]. In brief, XC-HMC is a strategy to reduce the adverse effects of momentum flipping on sampling autocorrelation times in GHMC. GHMC uses Langevin integration of Hamiltonian dynamics as a proposal mechanism, and accepts or rejects each proposal according to a Metropolis criterion. Whenever the Metropolis test fails, the proposal is rejected and the momentum must be reversed; this is necessary to maintain detailed balance[3,49], but can lead to extremely large autocorrelation times when the acceptance rate is not sufficiently close to 100% (see [43] for empirical examples and further discussion). In “extra-chance” HMC [41,43], rather than immediately flipping the momentum whenever the Metropolis criterion fails, the proposal trajectory is instead extended to generate a new proposal, and another (suitably modified) Metropolis criterion is checked. For suitable choices of parameters (length of trajectory proposal, timestep, number of extra chances, length of “extra-chance” trajectories), this strategy can virtually eliminate the effect of momentum flipping, at the cost of increasing the average length of proposal trajectories.

Our initial experiments (on larger systems than reported here) suggested that the cost of collecting uncorrelated samples using GHMC without “extra-chances” was prohibitive, since we needed to make the timestep extremely small (around 0.1–0.25 fs) to keep the acceptance rate sufficiently near 100% that the effect of momentum flipping was acceptable. Instead, we equilibrated for approximately 1 ns (10^5^ XC-HMC iterations, 10 steps per XC-HMC proposal trajectory, 15 extra-chance trajectories per iteration, 1 fs per timestep) from an energy-minimized starting structure. We then saved one sample **x**^(*i*)^ every 10^4^ XC-HMC iterations afterwards.

To draw an i.i.d. sample from *π*(**x**, **v**), we draw a sample **x** uniformly from *π*^cache^, and then sample **v** from the equilibrium distribution of velocities conditioned on **x**, **v ∼** *π* (**x***|***v**). (In the presence of holonomic constraints, the velocity distribution is not independent of the configuration distribution. For example, if bond lengths involving hydrogen atoms are constrained, the velocity of a hydrogen minus the velocity of the bonded heavy atom cannot have any component parallel to the bond.)

To draw an i.i.d. sample from *ρ*, we draw an i.i.d. sample from *π* and then simulate Langevin dynamics for a large number of steps. We tested using 1000 steps, (1/*γ*)/Δ*t* steps, and (2/*γ*)/Δ*t* steps.

### 6.4. Computing shadow work for symmetric Strang splittings

Here, we demonstrate how to compute the appropriate shadow work for a given discrete-timestep Langevin integration scheme. We note that the log-ratio of forward and reverse conditional path probabilities under specific time-discretizations of Langevin dynamics has been calculated in many prior works, such as [10,42,50]. While there has been a great deal of confusion in the literature about how nonequilibrium work should be computed [51], fortunately, there is an unambiguous mechanical (if tedious) approach to the computation of the appropriate work-like quantity.

Assemble the sequence of all steps and substeps of an integrator cycle into a trajectory *Z*. For example, for `OVRVO`, we have

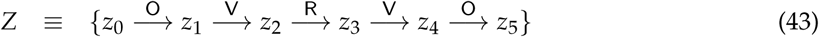

where *z_n_* ≡ (**x***_n_*, **v***_n_*) are phase-space points. Let 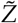 denote the time-reversal of all substeps of *Z* with all velocities negated,

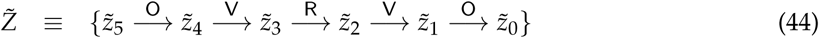

where 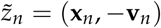 denotes negating the velocity of phase-space point *z_n_*.

To compute the reduced, unitless shadow work *w*[*Z*], we use the definition of work that satisfies the Crooks fluctuation theorem for starting with a sample from the target equilibrium density *π*(*z*_0_) and taking one integrator cycle step (eqn 4 of [1]):

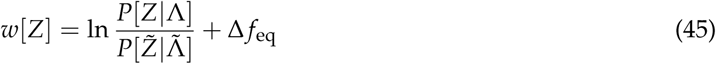

where the target equilibrium density is *π*(*z*) = *e* ^*f −h*(*z*)^, *f* is a log normalizing constant (dimensionless free energy), *h*(*z*) ≡ *u*(**x**) + *t*(**v**) is the reduced Hamiltonian, *u*(**x**) the reduced potential, and *t*(**v**) the reduced kinetic energy [52]. The quantity 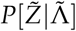 denotes the probability of generating the time-reversed trajectory 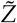 by starting with 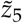 drawn from the target equilibrium density 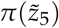 (and *not* the nonequilibrium steady state) and applying the reverse sequence of integrator operations 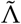, which is identical to the forward sequence of integrator operations Λ because the integrators we consider here are symmetric. Since the Hamiltonian is time-independent, the free energy change Δ *f*_eq_ = 0, and this simplifies to

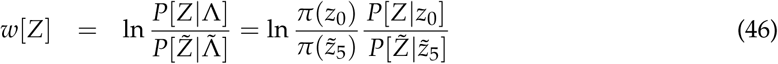

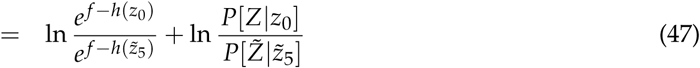

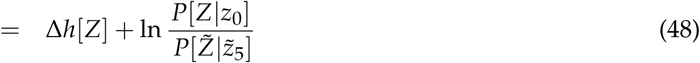

Computation of the shadow work then proceeds by simple mechanical algebra by computing the log ratio of conditional path probabilities in the last term.

For the family of integrators considered here (symmetric Strang splittings of the propagator, composed of `R, V`, and `O` steps), the shadow work has an especially simple form:

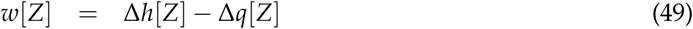

where Δ*h* is the total change in reduced Hamiltonian, and Δ*q* is the total change in reduced heat across each of the `O` substeps. We note that accumulation of the shadow work during integration requires no extra force evaluations, and simply requires knowledge of the potential energy at the beginning and end of the integrator cycle as well as the changes in kinetic energy for each `O` substep.

For `OVRVO`, this is

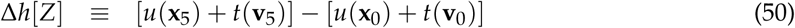

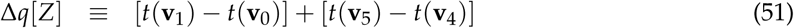

We illustrate how to arrive at this result in detail for `OVRVO` below.

### 6.5. *Computation of shadow work for* OVRVO

For `OVRVO`, we can represent the forward and reverse of a single integrator cycle diagrammatically as

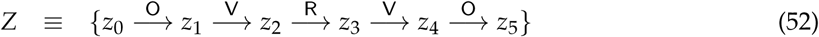

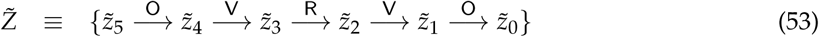

where 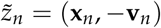 denotes negating the velocity of phase-space point *z_n_*.

To compute the conditional path probability *P*[*Z*|*z*_0_], we write a transition probability density kernel for each substep:

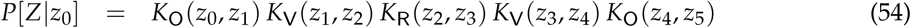

We can write the log ratio of conditional path probabilities as

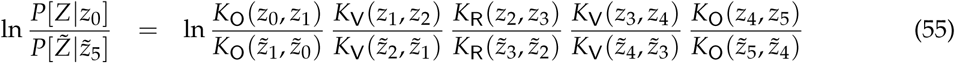

The probability kernels *K*_V_ and *K*_R_ are both deterministic, so as long as we are considering a trajectory *Z* and its time-reverse 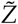 generated by a symmetric integrator splitting, the ratios involving these kernels are unity.

To compute the ratios involving *K*_O_ kernels, we note that *K*_O_(*z*_0_, *z*_1_) perturbs the velocity according to the update equation

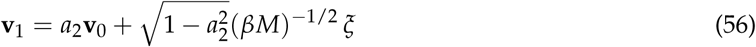

where *ξ* is a random variate drawn from the unit normal density, which allows us to solve for the random variate required to propagate from *z*_0_ to *z*_1_,

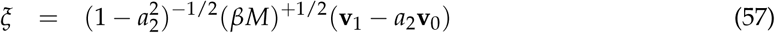

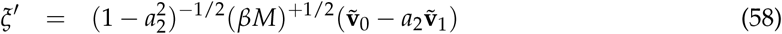

where the probability density is given by

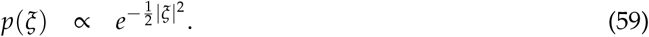

We can then rewrite the log ratio of `O` kernels as

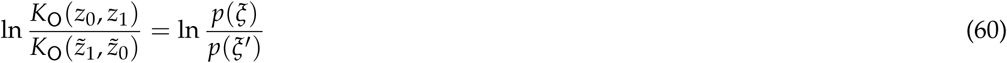

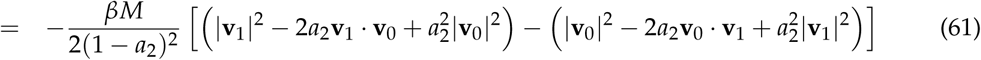

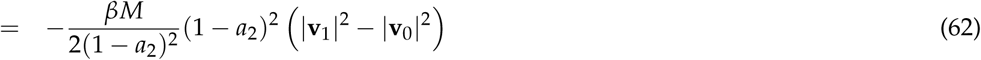

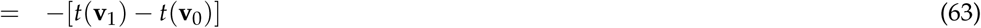

Combining this with (49), this provides the overall work as

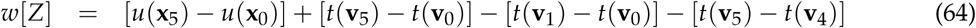

### 6.6. Variance-controlled adaptive estimator for KL divergence

As noted in Section 3.6, the nested Monte Carlo estimator we use as an expensive, but in principle asymptotically exact, estimate of the KL divergence requires converging a separate exponential average ln〈*e^−w^*〉_**x,v**;Λ_ for every sample (**x**, **v**) ∼ *ρ* or *ω*. It is obviously impossible to compute this exactly; any practical approach employing finite computational resources can only estimate this quantity to some finite statistical precision, and even then, the logarithm will induce some bias in the computed estimate for finite sample sizes. Here, we take the approach of setting a sensible target statistical error for this inner estimate, arbitrarily selecting 0.01, since we would like to resolve features in 𝒟_KL_ larger than this magnitude. Notably, the difficulty in achieving this threshold increases exponentially as the width of the sampled distribution *p*(*w*) increases with increasing timestep Δ*t*.

To determine the number of inner-loop samples required to meet this statistical error threshold, we periodically compute an estimate of 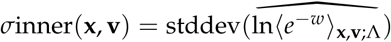 from the available samples using standard first-order Taylor series propagation of uncertainty. We compare the estimated standard deviation with the user-defined threshold, and continue to draw samples until we meet the threshold or exceed the sample budget of 5 × 10^4^ samples^8^. The scaling of computational effort with Δ*t* is shown in Figure A4.

Choosing the inner-loop threshold is subtle. If the inner-loop threshold is chosen too large, then the resulting estimate will be very biased (the whole procedure only becomes unbiased in the limit that *σ*inner → 0). If the threshold is chosen too small, then the computational effort becomes prohibitive. Controlling the *precision* of the inner-loop estimates should also be expected to control their *bias*, since the bias of the inner-loop estimates is approximately 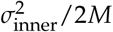 (see Section II.B, eqn. 8 in [40]), in the direction of *under-estimating* the 𝒟_KL_.

To compute and report uncertainty in Figure 7, we use bootstrap resampling, rather than Taylor propagation. The data for each condition is a jagged array of sampled work values, where each row represents an “outer-loop sample” (*i.e*., a different initial condition (**x**, **v**) sampled from *ρ* (or *ω*)), and the length of each row is variable, reflecting the number of “inner-loop” samples required to estimate ln [*ρ* (**x**, **v**)/*π*(**x**, **v**)] (or ln [*ω*(**x**, **v**)/*π*(**x**, **v**)]) to the desired precision. To generate a single bootstrap sample, we resample first the rows uniformly with replacement, and then, within each row, resample the columns uniformly with replacement. The error bands for the “exact” estimator in Figure 7 are computed from 100 bootstrap samples per condition.

## Appendix A Statistics of shadow work distributions

The exact expression for the KL divergence used for validation in Section 3.6 requires estimating the expectation of *e^−w^* averaged over *p*(*w*). Work distributions for various integrators and timesteps are plotted in Figure A1, and appear to be approximately Gaussian, as can be seen by comparison with Gaussian fits (solid lines). As expected, the width of the work distribution increases with increasing timestep, which can be seen more clearly in Figure A2, which plots the standard deviation of the work distribution as a function of timestep.

**Figure A1.**
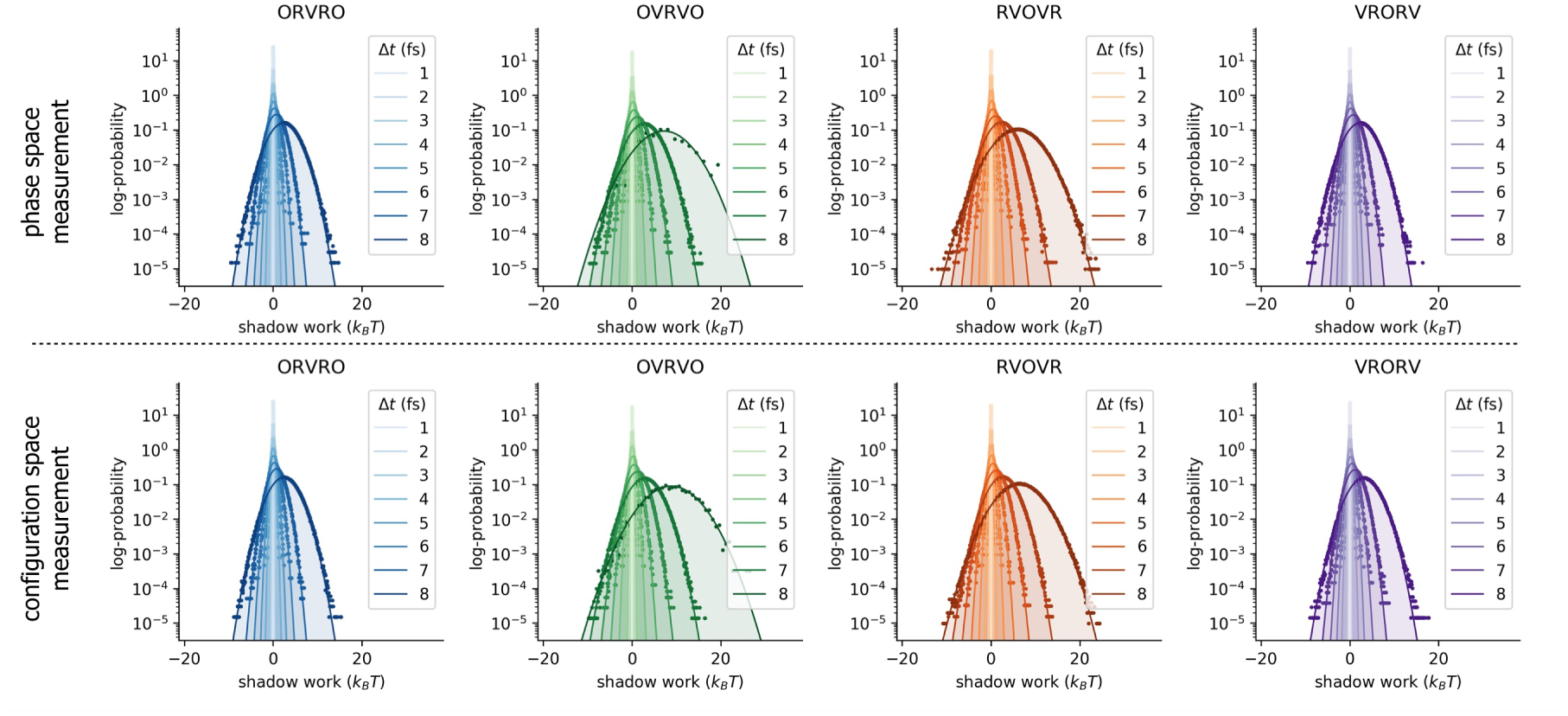
Shadow work distributions for the water cluster are approximately Gaussian for all integrators examined. In all panels, solid lines and shaded regions denote Gaussian fits, while dots denote histogram estimates. The **top row** depicts work distributions where initial conditions are sampled from the nonequilibrium steady-state induced by the corresponding integrator and timestep [(**x**, **v**) ~ *ρ*]; these shadow work values are used to measure phase-space error in the near-equilibrium estimates of 𝒟_KL_. The **bottom row** depicts work distributions where initial conditions are sampled from the *w* ensemble [**x** ~ *ρ***_x_**, **v** ~ *π*(**v**|**x**)]; these work values are used to estimate configuration-space error in the near-equilibrium estimates of 𝒟_KL_.

**Figure A2.**
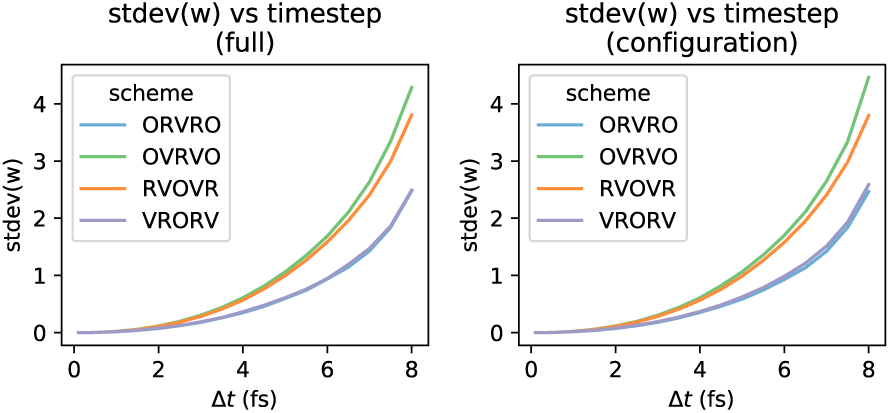
Standard deviation of water cluster shadow work distribution grows with time step. The left panel summarizes the top row of Figure A1 (shadow work distributions for trajectories initialized in *π*), and the right panel summarizes the bottom row of Figure A1 (trajectories initialized in *ω*).

While the near-equilibrium estimate will find the difficulty of reaching estimates of a given statistical precision grows linearly with the variance in *p*(*w*), the exact estimator must converge expectations of the *exponentiated* shadow work, which becomes exponentially difficult with increasing variance. This effect is illustrated in Figure A3, where we plot *e^−w^* along with the Gaussian fits to these work distributions.

This is reflected in the computational effort required by the variance-controlled estimator, illustrated in Figure A4.

**Figure A3.**
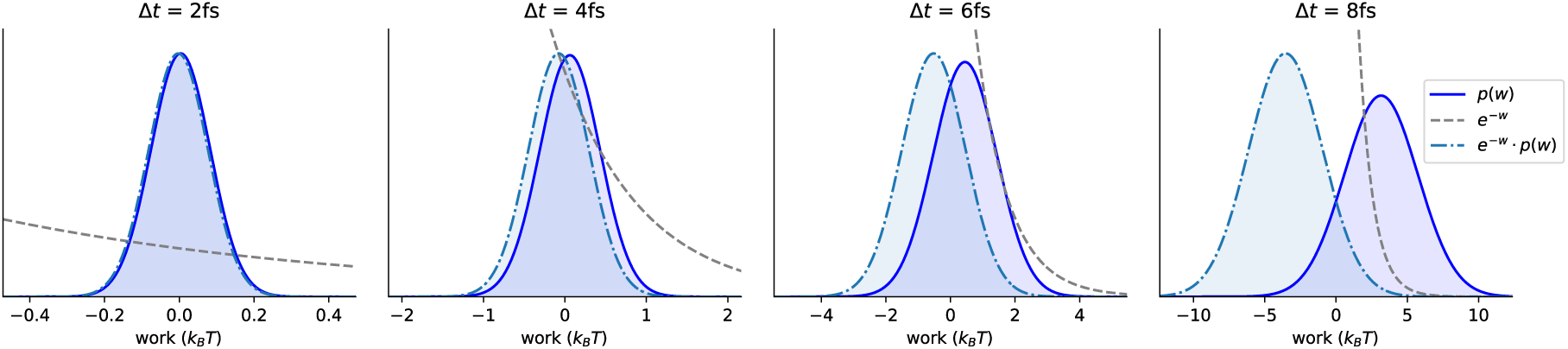
Exponential averages with respect to the shadow work distribution become increasingly difficult with increasing timestep. It becomes increasingly difficult to estimate the expectation of *e*^−*w*^ with respect to Gaussian fits to work distributions *p*(*w*) for the water cluster, as the timestep Δ*t* increases. The four panels increase in Δ*t* from left to right: note the changing *x*-axis scales. The **solid line** is the shadow work distribution *p*(*w*) measured at each timestep. The **dashed line** is *e^−w^*. The **dash-dotted line** is *e*^−*w*^ · *p*(*w*).

**Figure A4.**
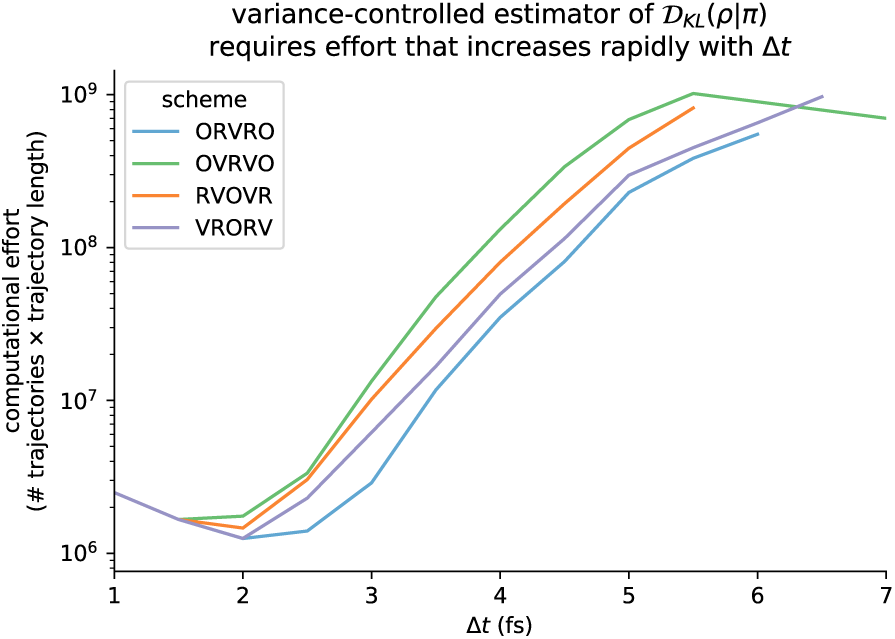
The computational effort required to reach a fixed uncertainty threshold depends sharply on Δ*t*. The total computational effort is determined by the total number of trajectories sampled in that condition (*i.e*. 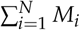), multiplied by the length of each trajectory. Note that the curves are not monotonic, since the number of required trajectories increases superlinearly with Δ*t*, but the number of timesteps in each trajectory decreases linearly with Δ*t*.

## Appendix B Log-scale plots

To better visualize agreement between near-eq and nested estimates at small timesteps, we also show a version of Figure 7 with a log-scale *y*-axis in Figure A5.

**Figure A5.**
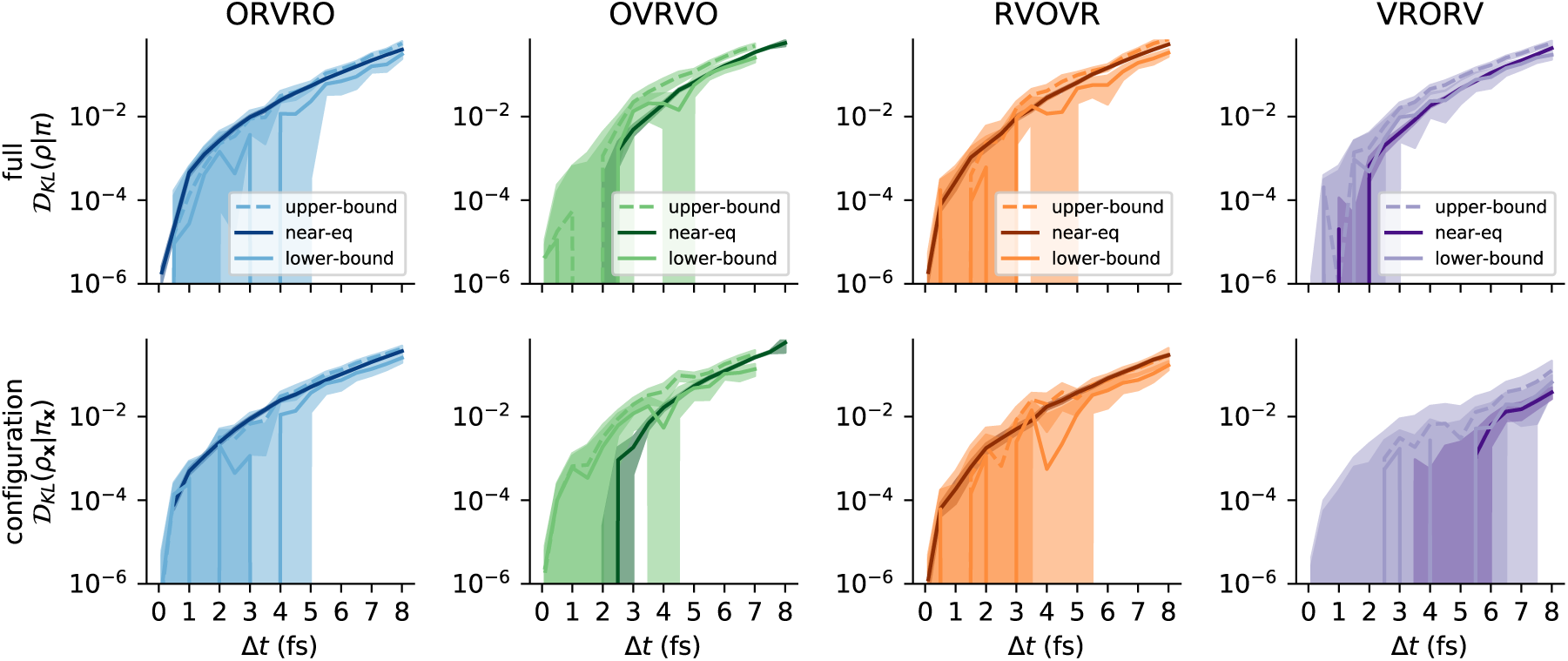
Agreement between near-equilibrium and reference methods may be better visualized at small timesteps on a log-scale. This is a repeat of Figure 7, but with a log-scale *y*-axis. Vertical artifacts are caused by the estimate becoming transiently negative, which cannot be represented on the logarithmic scale.

## Appendix C Further comments on the collision rate

If we are interested only in configuration-sampling, the collision rate is a free parameter. However, the collision rate affects the sampled kinetics. Depending on the application, we may choose a value of the collision rate that reproduces kinetic observables such as diffusion constants, or we may choose a value of the collision rate that minimizes correlation times. In the over-damped limit (large *γ*) conformational search is significantly hampered, as the trajectories become diffusive. In the under-damped limit (small *γ*) trajectories are not random walks, but mixing between energy levels is slower. (At the extreme, *γ* = 0, Langevin dynamics reduces to Hamiltonian dynamics, and explores a single constant-energy ensemble.)

We do not provide any guidance here on the choice of the collision rate, but we investigated how sensitive the measured 𝒟_KL_ is to the choice of *γ* for each integrator. To supplement the figure in the main text (Figure 6), we plot the same information again, but grouped by *γ* rather than by scheme in Figure A6.

**Figure A6.**
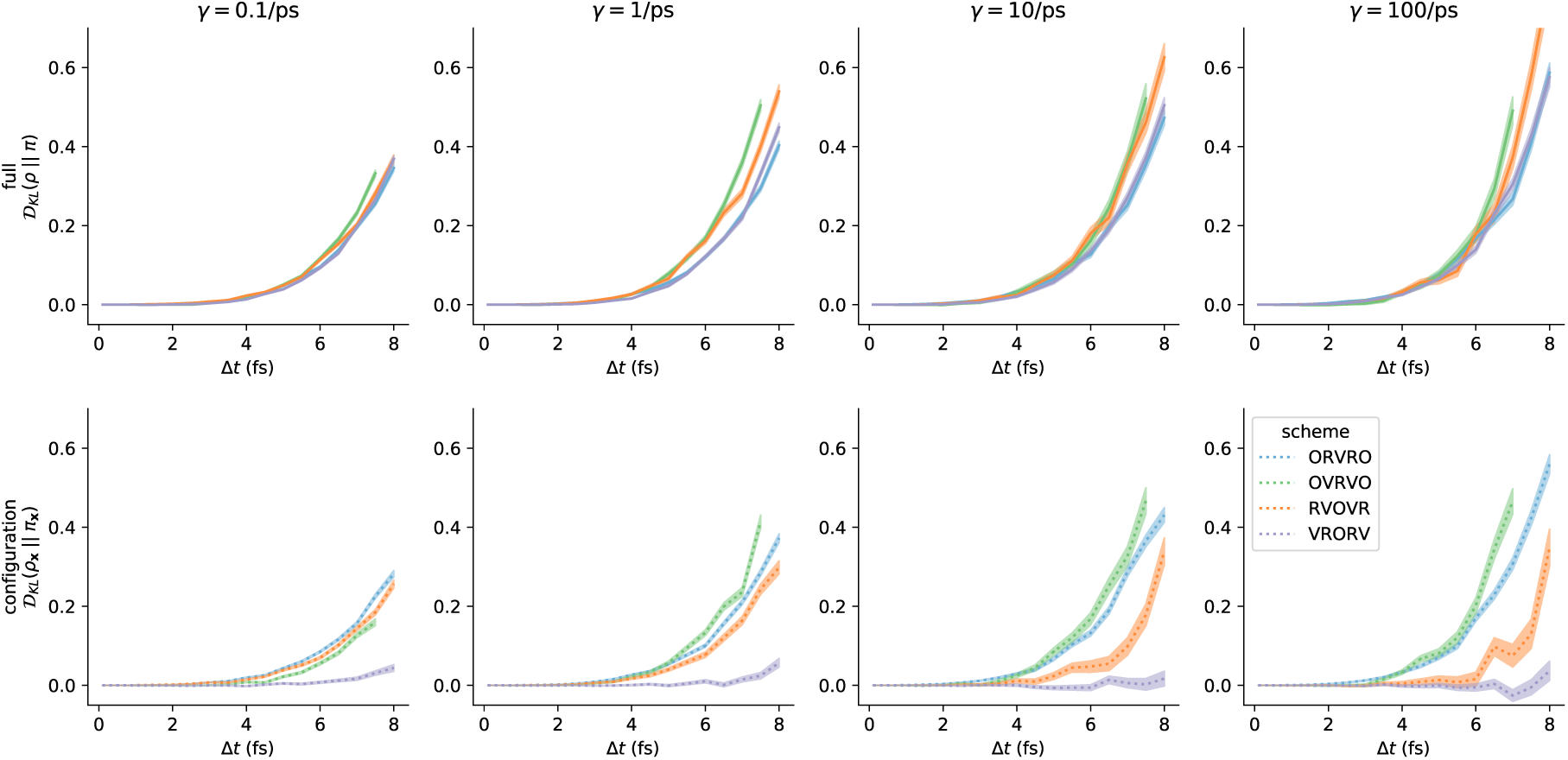
Integrator accuracy can be compared at different collision rates. Here we have plotted the same results as in Figure 6 but grouped by the collision rate *γ* rather than by scheme. Note that the ordering over schemes induced by configuration-space error can vary as a function of collision rate—*i.e*. at the lowest measured *γ*, the ordering from lowest to highest error is (1) `VRORV`, (2) `OVRVO`, (3) `RVOVR`, (4) `ORVRO`, but at the highest measured *γ*, the ordering is different.

As noted above, the accuracy of the proposed estimator hinges on the assumption that the system has been driven into steady-state during the finite-length protocol. To validate that protocols of length 2000 steps were sufficient, we compare results for protocols that are 1000 steps long and 2000 steps long, and note that they produce very similar estimates for timestep-dependent error for the range of collision rates examined. The results reported in Figure 6 and Figure A6 were performed using a protocol of length *T* = 2000 steps, two times longer than the protocols used in Figure 5. We also computed near-equilibrium estimates over the same range of collision rates using a shorter protocol of length of *T* = 1000 steps Figure A7. Note that, for computational convenience, we have used substantially fewer protocol samples to compute the estimates in Figure A7 than were used to compute the results reported in the main text (10 000 samples, rather than 50 000 samples).

**Figure A7.**
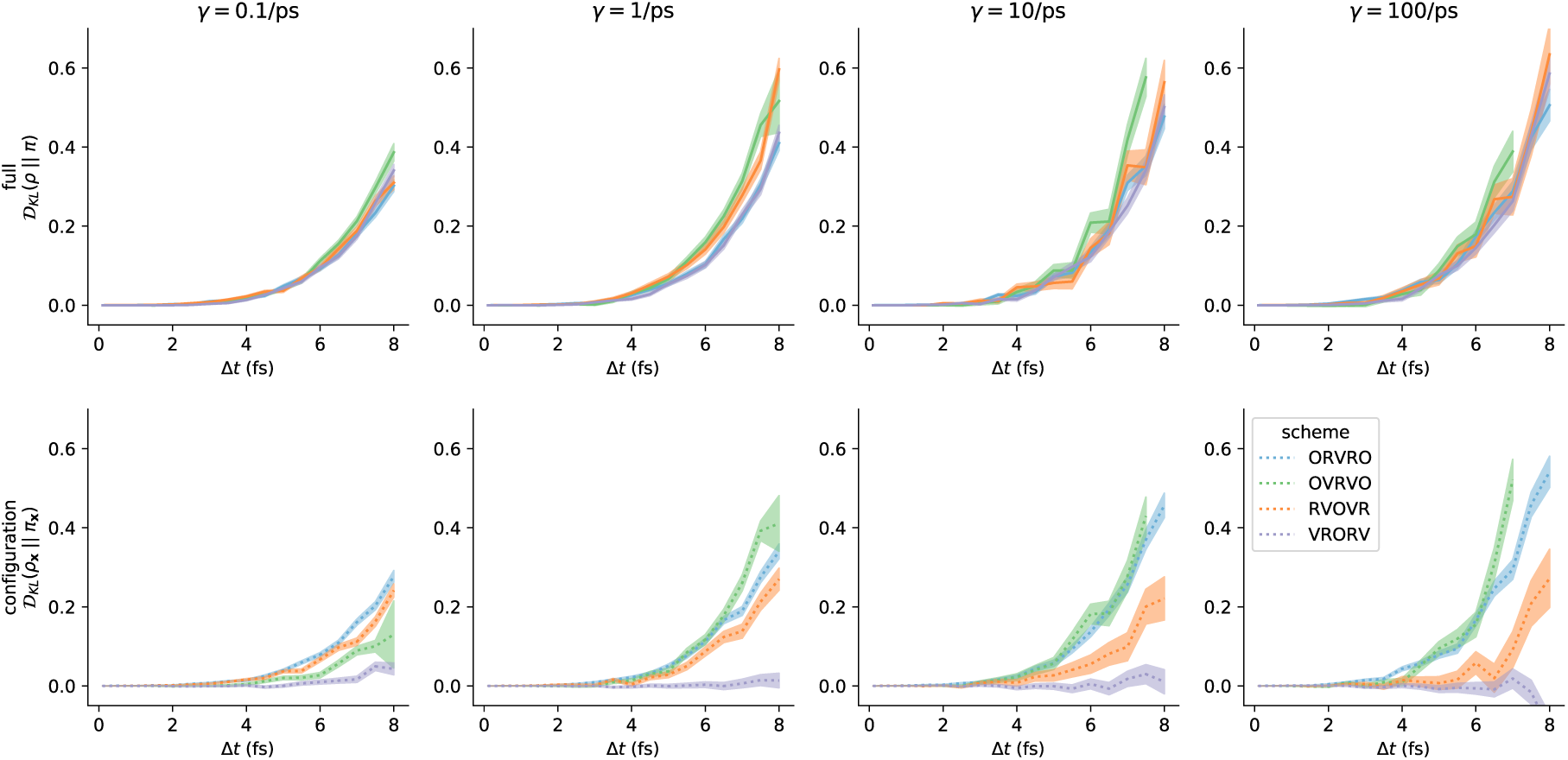
Similar results are obtained for 1000-step protocols as for 2000-step protocols. For the same wide array of conditions reported in Figure A6, near-equilibrium estimates generated using 5x fewer samples of 2x shorter protocols are broadly consistent with the results obtained by more exhaustive sampling.

## Acknowledgments

The authors gratefully acknowledge the helpful suggestions of Grant Rotskoff (ORCID: 0000-0002-7772-5179) for his input on nonequilibrium shadow work measurement schemes; Charles Matthews (ORCID: 0000-0001-9545-1346) and Brian Radak (ORCID: 0000-0001-8628-5972) for discussions on geodesic integrators; Jason Wagoner (ORCID: 0000-0003-2359-9696) for discussions about Metropolized Langevin integrators; and members of the Chodera laboratory for input on various aspects of the implementation and theory. We thank Rodrigo Guerra (ORCID: 0000-0003-3082-0248) for helpful comments on continuous barostats. The simulations were made possible by the High Performance Computing Group at MSKCC. BJL was supported by EPSRC grant no. EP/P006175/1 and the Alan Turing Institute. JDC acknowledges support from the Sloan Kettering Institute and NIH grant P30 CA008748 and NIH grant R01 GM121505. JF acknowledges support from NSF grant CHE 1738979. DAS acknowledges support from a Natural Sciences and Engineering Research Council of Canada (NSERC) Discovery Grant and the Canada Research Chairs program. We are especially grateful to Peter Eastman (ORCID: 0000-0002-9566-9684) for implementing the CustomIntegrator facility within OpenMM that greatly simplifies implementation of the integrators considered here, and for helpful feedback on the manuscript. Finally, we thank the three anonymous reviewers for their many thoughtful comments that greatly improved the manuscript, for identifying relevant prior work we had missed, and for suggesting additional experiments to isolate the effect of the collision rate.

## Footnotes

1 The concept of a *shadow Hamiltonian* has been used to embed this density in a canonical density context, but the shadow Hamiltonian cannot be directly computed, though some approaches to approximate it via expansion (generally requiring higher-order derivatives than gradients) have been proposed—see [21], Chapter 3 of [4], and references therein.

2 The subscripts 1/4,1/2, etc., have no relation to intermediate physical times, and are used solely to denote the sequence of intermediate computational steps.

3 In the limit that Δ*t* → 0, this relation is exact, since 𝒟_KL_(*ρ*||*π*) → 0 and both 〈*w*_shad_〉_*π*;∧_ → 0 and 〈*w*_shad_〉_*ρ*;∧_ → 0.

4 Spatial discretization or density estimation are infeasible, due to curse of dimensionality. There are direct estimators of the KL divergence based on Euclidean nearest-neighbor distances that perform well in some high-dimensional settings (*e.g*., [37]), but Euclidean distance is an unsuitable metric on molecular configurations.

5 Note that applying this methodology to non-symmetric integrators (where the sequence of operations for the integrator its time-reverse are not identical) would require modifications to this scheme, as well as the manner in which shadow work is computed.

6 Note that when we approximate this bound with a finite number of samples, we will underestimate it for the reasons mentioned in Section 3.6.1. However, since we are pooling all work samples, the magnitude of this underestimate should be much smaller than for the inner-loop underestimates in Section 3.6.1 above, and we expect the magnitude of this bias to be negligible compared with the effect of invoking Jensen’s inequality.

7 In [23], the error in phase space was measured for an ensemble of constant-volume Langevin trajectories with initial conditions drawn from the isothermal-isobaric ensemble.

8 This limited the maximum CPU time spent collecting an “outer-loop” sample to approximately 2 hours—conditions where this limit was met are colored grey in the lower panel of Figure 7.

